# Intrinsic Gating Behavior of Voltage-Gated Sodium Channels Predetermines Regulation by Auxiliary β-subunits

**DOI:** 10.1101/2021.02.25.432706

**Authors:** Niklas Brake, Adamo S Mancino, Yuhao Yan, Takushi Shimomura, Heika Silveira, Yoshihiro Kubo, Anmar Khadra, Derek Bowie

**Author notes:** These authors contributed equally to this work (Niklas Brake, Adamo S Mancino). Senior author (Anmar Khadra, Derek Bowie).

## Abstract

Voltage-gated sodium (Nav) channels mediate rapid millisecond electrical signaling in excitable cells. Auxiliary subunits, β1-β4, are thought to regulate Nav channel function through covalent and/or polar interactions with the channel’ s voltage-sensing domains. How these interactions translate into the diverse and variable regulatory effects of β-subunits remains unclear. Here, we find that the intrinsic movement order of the voltage-sensing domains during channel gating is unexpectedly variable across Nav channel isoforms. This movement order dictates the channel’ s propensity for closed-state inactivation, which in turn modulates the actions of β1 and β3. We show that the differential regulation of skeletal muscle, cardiac, and neuronal Nav channels is explained by their variable levels of closed-state inactivation. Together, this study provides a unified mechanism for the regulation of all Nav channel isoforms by β1 and β3, which explains how the fixed structural interactions of auxiliary subunits can paradoxically exert variable effects on channel function.

## Main Text

### Introduction

Almost all ligand-and voltage-gated ion channels expressed in excitable tissue assemble as a signaling complex consisting of pore-forming and auxiliary subunits (Dolphin, 2018; Edelheit et al., 2009; Gonzalez-Perez and Lingle, 2019; Hull and Isom, 2018; Twomey et al., 2019). For example, the voltage-gated sodium (Nav) channel complex assembles from one of nine possible pore-forming α-subunits (Nav1.1-Nav1.9) with 1-2 auxiliary β-subunits (β1 to β4) that together shape the rapid and varied kinetics of action potentials in brain, heart and skeletal muscle tissue (Catterall et al., 2005; Hull and Isom, 2018). Individual Nav channel α-subunits form a tetrameric structure around a central Na^+^-selective pore from four non-identical domains (DI-DIV). Each domain contains 6 transmembrane segments (S1-S6) that are either responsible for voltage detection (S1-S4) or formation of the pore structure (S5-S6) (Ahern et al., 2016; Jiang et al., 2020; Pan et al., 2018). Biochemical and electrophysiology studies as well as recent full-length cryo-EM Nav structures have revealed that these domains are selectively targeted by β-subunits: β1 and β3 subunits form polar interactions with DIII or DIV (Hsu et al., 2017; Hull and Isom, 2018; Pan et al., 2018; Yan et al., 2017) whereas β2 and β4 establish covalent links with DI and/or DII (Das et al., 2016; Hull and Isom, 2018; Isom et al., 1995, 1992; Shen et al., 2019). Together, these findings establish a common structural view of the interactions between α-and β-subunits.

Despite this, the effects of β1 and β3 on channel function remain undetermined. Although numerous electrophysiological studies have characterized the effects of β1, in the two most heavily studied Nav channels Nav1.4 and Nav1.5 – β1 has been shown to produce large (Bendahhou et al., 1995; Zhu et al., 2017), moderate (Malhotra et al., 2001; Nuss et al., 1995), or even no changes (Ferrera and Moran, 2006; Nuss et al., 1995) to channel gating. This variability, even within a single Nav channel isoform, demonstrates that the emerging structural view of β-subunits has been, so far, insufficient to explain their effects on channel function.

Here, we attempted to reconcile this apparent disconnect between structural and electrophysiological studies. Because β-subunits have been shown to alter voltage sensor movements to produce their functional changes in Nav1.5 channels (Zhu et al., 2017), we began by investigating the contributions of each voltage sensor to the gating of Nav1.5e, a neonatal form of Nav1.5 that is expressed in the brain (Wang et al., 2017). Using voltage-sensor neutralization experiments, voltage-clamp fluorometry (VCF), and kinetic modelling, we find that the functional contributions of each voltage sensor are not fixed. Instead, we demonstrate that the sequence of voltage sensor movements is variable, which in turn modulates the contributions of each voltage sensor to channel activation and inactivation. We further show that this mechanism determines the functional consequences of β1 and β3 association with cardiac Nav1.5 channels, skeletal muscle Nav1.4 channels, and neuronal Nav1.6 channels. Finally, through an analysis of previously published studies, we find that this mechanism successfully explains the variable regulation of all Nav channel isoforms by β1. We conclude that β1 and β3 impose a defined structural influence on all Nav channels and that their variable allosteric effects are determined by the intrinsic dynamics of the channel’ s voltage sensors.

## Results

### Voltage Sensor Charge Neutralization Identifies the Dominant Role of DI in Channel Activation

Voltage-detection in Nav channels is mediated by gating charges (Arg or Lys residues) on each S4 segment (Fig. 1A) which together promote movement in individual S3-S4 voltage sensors following changes in the membrane electric field. To study the contribution of individual S3-S4 voltage sensors to channel gating, the first three gating charges in each S4 segment of the mouse Nav1.5e channel were mutated to glutamines to generate four charge-neutralized (CN) mutant channels: DI-CN, DII-CN, DIII-CN, DIV-CN (Fig. 1A). We reasoned that neutralization of individual voltage sensors would render the domain insensitive to changes in the membrane potential and, as such, would inform us about the biophysical properties of Nav1.5e, as reported for other voltage-gated sodium and potassium channels (Bao et al., 1999; Capes et al., 2013, 2012; Gagnon and Bezanilla, 2009; Sheets et al., 1999). Nav1.5e channels and the charge-neutralized mutants were characterized in HEK-293T cells transiently transfected with wildtype and mutant cDNAs. To assess channel activation, macroscopic Na^+^ currents were recorded by applying depolarizing voltage steps in increments of +5 mV (Range, −110 and 55 mV), each from a holding potential of −130 mV (−100 mV for wildtype channels) (Fig. 1B). The voltage-dependence of channel activation for wildtype and mutant channels was then determined by fitting the peak conductance (G/V) with a Boltzmann function (Fig. 1C).

**Figure 1.**
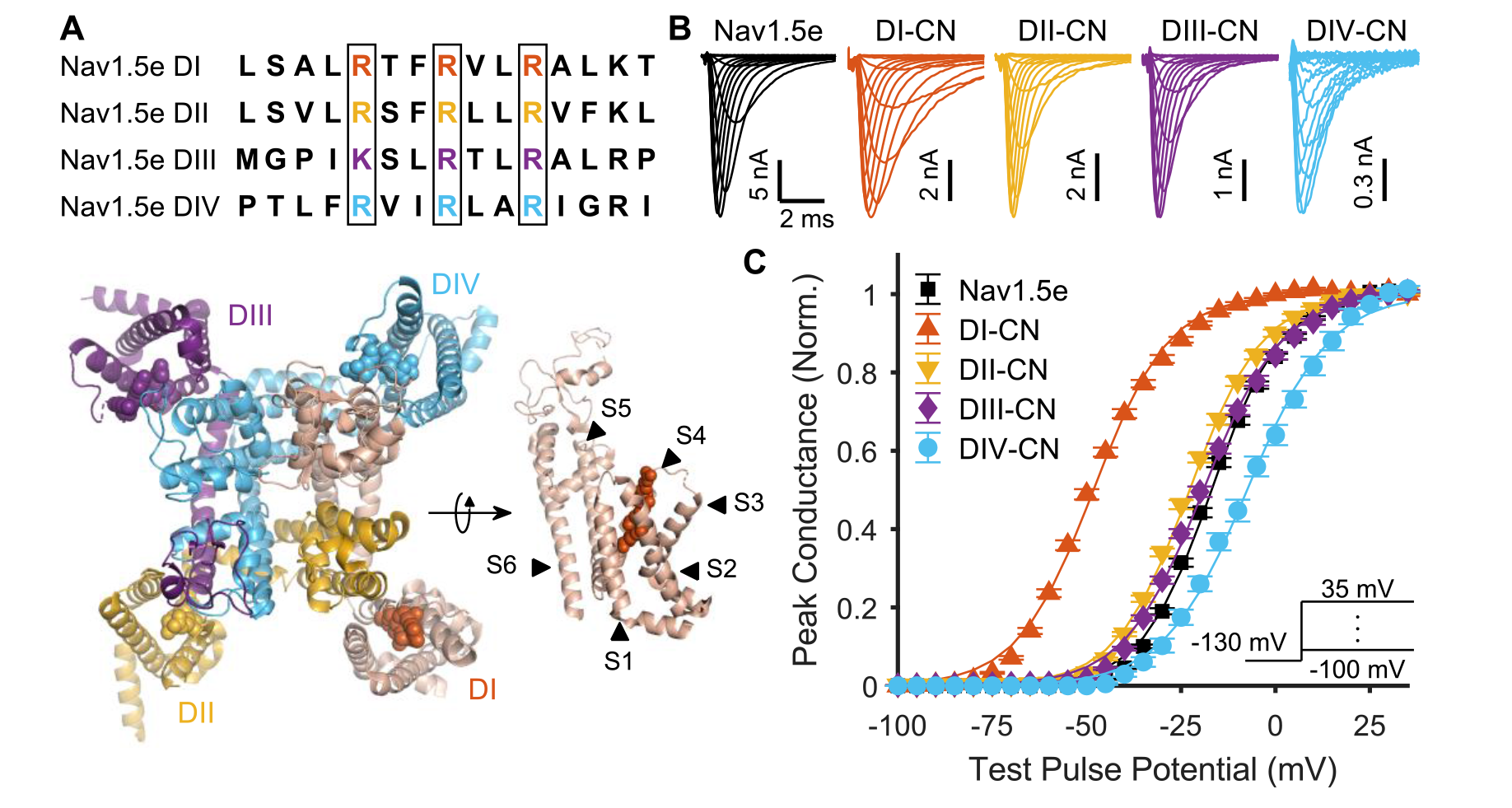
Impact of neutralizing voltage-sensing domains on channel activation. (**A**) Top: Sequence alignment of the S4 voltage-sensing helices across domains I-IV of mNav1.5e. Outlined in black are the gating charges that were neutralized to glutamine in the charge-neutralized (CN) mutants. Bottom: top-down view of the structure of Nav1.5 from (Jiang et al., 2020). To the right is a side view of domain I with the various transmembrane α helices annotated. Gating charges corresponding to those outlined in black boxes in the top of the panel are annotated as balls in the structure. (**B**) Representative traces of ionic currents corresponding to wild-type (WT) Nav1.5e (cell 20170418c2) and mutant channels (DI-CN, cell 20180307c1; DII-CN, cell 20180316c1; DIII-CN, cell 20180712c3; DIV-CN, cell 20190318c3) in response to depolarizing voltage steps ranging from −110 to 35 mV, following a holding potential of −130 mV (−100 mV for WT), recorded in HEK-293T cells. A scheme of this voltage step protocol is displayed in the inset of panel **C**. (**C**) Normalized peak conductance (GV) of wild-type and mutant channels. Inset: voltage step protocol used to assess channel activation. Solid lines are fitted Boltzmann curves (Table S1).

Charge neutralization led to statistically significant (see methods) shifts in the activation profile of all mutant Nav1.5e channels, with DI charge neutralization producing the greatest impact (Fig. 1C). Fits of peak G/V relationships estimated the voltage for half-maximal activation (V_1/2_) to be −16.8 ± 0.45 mV (n = 51) for wildtype channels compared to −48.1 ± 0.55 mV (n = 24) for DI-CN (Fig. 1C, Table S1), representing a 30 mV hyperpolarizing shift in channel activation. The slope factor (k) was similar for wildtype (k = 9.0 ± 0.13 mV, n = 51) and DI-CN channels (k = 9.9 ± 0.13 mV, n = 24). In contrast, charge neutralization of DII and DIII had a more modest effect on channel activation with V_1/2_ values of −22.6 ± 0.54 mV (n = 26) and −19.0 ± 0.58 mV (n = 23), respectively (Fig. 1C), corresponding to hyperpolarizing shifts in activation of about 6 and 2 mV compared to wildtype Nav1.5e (Table S1). Finally, charge neutralization of DIV had the opposite effect on channel activation, shifting the V_1/2_ value to −7.4 ± 1.26 mV (n = 14) (Fig. 1C), representing a 10 mV *depolarizing* shift in channel activation compared to wildtype Nav1.5e (Table S1). Similar relative shifts in channel activation were observed when DI through to DIV voltage sensors were charge neutralized in the adult form of Nav1.5 (i.e. mH1), demonstrating that our observations are not specific to the neonatal form of Nav1.5 (Fig. S1). Our observations on Nav1.5 are comparable with previous findings on skeletal muscle Nav1.4 channels, with the exception of DIV-CN which did not depolarize activation in Nav1.4 (Capes et al., 2013).

### DI Movement Is the Rate-Limiting Step for Pore Opening

The canonical Nav channel gating model postulates that DI-III movements are necessary for pore opening and DIV movement is sufficient and rate-limiting for inactivation (Ahern et al., 2016). To understand why neutralizing DIV affected activation (Fig. 1C), we attempted to explain our Nav1.5e data using a mathematical implementation of this gating model (Fig. 2A) (Capes et al., 2013). The model successfully captured the wildtype Nav1.5e data after the rate constants were re-parametrized using a custom-made evolutionary-type fitting algorithm (Fig. S2, Table S2) (see Methods).

**Figure 2.**
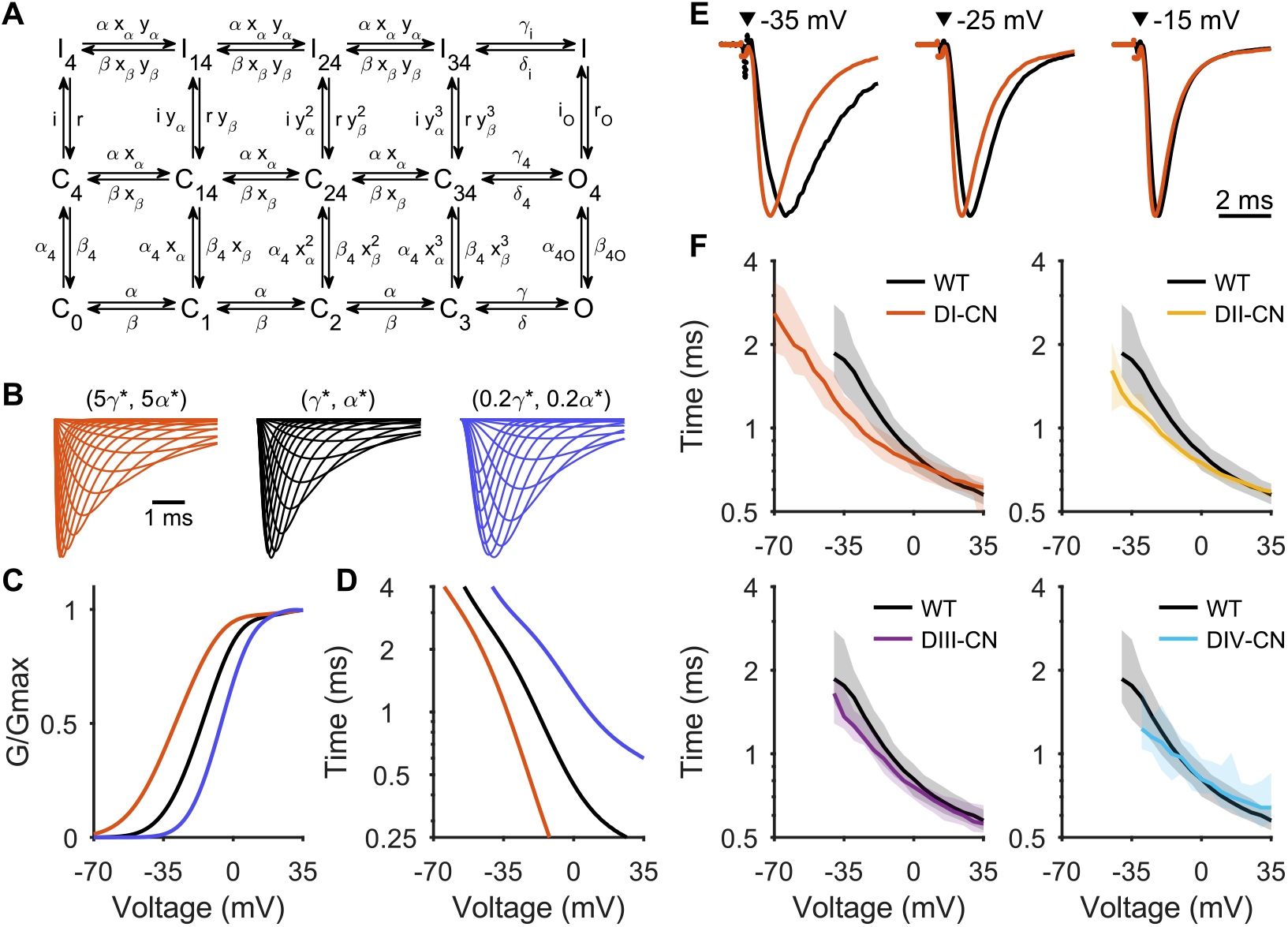
A model of Nav gating predicts changes in current rise time. (**A**)A re-parameterized kinetic model of Nav gating, adopted from Capes *et al*. (2013). Horizontal transitions from left to right represent the nonspecific movement of DI-III, followed by pore opening. Vertical transitions from bottom to top represent the movement of DIV followed by the movement of the inactivation gate. See Fig. S2 for a comparison between the model and data. (**B**) Middle: Representative traces of the WT Nav1.5 model response to the activation protocol. Right: γ* and α* have both been reduced by 80% to simulate a slower activation rate. Left: γ* and α* have both been increased by 400% to simulate a faster activation rate. See Fig. S3 for full sensitivity analysis. (**C**) GV plots calculated from the traces of corresponding colour in panel **B**. (**D**) Time to peak current calculated from the traces of corresponding colour in panel B. (**E**) Representative traces showing the current (normalized to peak amplitude) elicited by the indicated voltage steps for WT (black; cell 20170428c1) and DI-CN mutant channels (red; cell 20180307c1). (**F**) Time to peak current induced by voltage pulses ranging between −75 and 30 mV, starting from a holding potential of −130 mV, for Nav1.5e (black) and the four CN mutants. Line represents median while shading is 5-95% quantile interval. Time to peak was not calculated at voltages that did not elicit a current.

We hypothesized that DI-CN hyperpolarizes G/V because DI movement is normally rate-limiting for pore opening and that, conversely, DIV-CN might depolarize the G/V relationship by slowing the rate of pore opening. In the kinetic gating model, the rates that determine the transitions from the resting state to the open state are α and γ (Fig. 2A). To investigate how the voltage-dependence of the G/V relationship depends on these rates, we performed a sensitivity analysis with respect to parameters α* and γ* (Table S2). This analysis indicated that if domain neutralization accelerates or slows the rate of pore opening, the G/V relationship should be hyperpolarized or depolarized, respectively (Fig. S2E, F). For example, decreasing α and γ by 80% led to a depolarizing shift in the G/V relationship while increasing the rates by 400% led to a hyperpolarizing shift (Fig. 2B, C), consistent with our expectations.

Additionally, our simulations revealed that a faster or slower rate of pore opening should accelerate or slow the response risetime, respectively (Fig. 2B, D). This observation is in line with single-channel studies, which have reported that the time course of macroscopic currents is primarily determined by the latency to first opening (Aldrich et al., 1983). To validate our hypotheses about DI and DIV, we therefore measured the risetime to peak current of each mutant Nav channel (Fig. 2E, F). As anticipated, DI-CN channels displayed faster risetimes at hyperpolarized membrane potentials (Fig. 2F), suggesting that DI movement is normally rate-limiting for pore opening. The risetimes of DII-CN and DIII-CN mutant channels were likewise consistent with their more modest impacts on the V_1/2_ of channel activation (Fig. 1C; Fig. 2F). Unexpectedly, DIV-CN mutants did not display slower response risetimes relative to WT channels (Fig. 2F), suggesting that neutralizing DIV does not slow the rate of pore opening. We therefore concluded that DIV-CN must shift the G/V relationship through a separate mechanism. Since DIV is intrinsically involved in Nav1.5 inactivation (Jiang et al., 2020), we hypothesized that charge neutralization of DIV may affect the voltage-dependence of activation indirectly because of the interdependence of the two processes. As explained below, we examined this by measuring the impact of charge neutralization on steady-state inactivation.

### Voltage Sensors of All Domains Contribute to Steady-State Inactivation of Nav1.5e Channels

Steady-state inactivation (SSI) of wildtype and mutant Nav1.5e channels was determined by applying a 100 ms-long conditioning pulse (range, −160 to −5 mV) followed by a test pulse of −10 mV to elicit Na^+^ currents (Fig. 3A, B). SSI plots were then constructed by fitting the peak response at each test potential with a Boltzmann function (Fig. 3C). In agreement with previous work on Nav1.4 (Capes et al., 2013), neutralizing DIV had the largest effect on SSI, shifting the V_1/2_ from −82.0 ± 0.52 mV in wildtype channels to −131.1 ± 2.55 mV (n = 13) in DIV-CN mutants (Table S1). The slope factor for the inactivation curve was significantly flatter for DIV-CN mutants (k = −13.7 ± 1.46, n = 13) than for wildtype Nav1.5e channels (k = −7.2 ± 0.14, n = 49), indicating a lower sensitivity to membrane potential. Charge neutralization of DIII affected SSI in a manner similar to neutralizing DIV, hyperpolarizing the V_1/2_ of inactivation by 40 mV to −120.0 ± 1.09 mV (n = 23) and flattening the SSI slope factor to −14.4 ± 0.15 (n = 23). Although measurements of SSI for DI and DII-CN mutants differed from wildtype Nav1.5e, the shift was less than for DIII and DIV-CN mutants. The V_1/2_ of inactivation was estimated to be −100.9 ± 0.54 mV (n = 23) and −96.5 ± 0.82 mV (n = 25) for DI and DII-CN mutants, respectively (Fig. 3C, Table S1) corresponding approximately to 20 mV and 15 mV hyperpolarizing shifts. Similar relative shifts in SSI were also observed from charge-neutralized adult Nav1.5 channel mutants (Fig. S3), again demonstrating that our observations are not specific to the neonatal form of Nav1.5. However, these results were starkly different from past studies on Nav1.4 (Capes et al., 2013), where only DIV-CN mutants displayed altered SSI.

**Figure 3.**
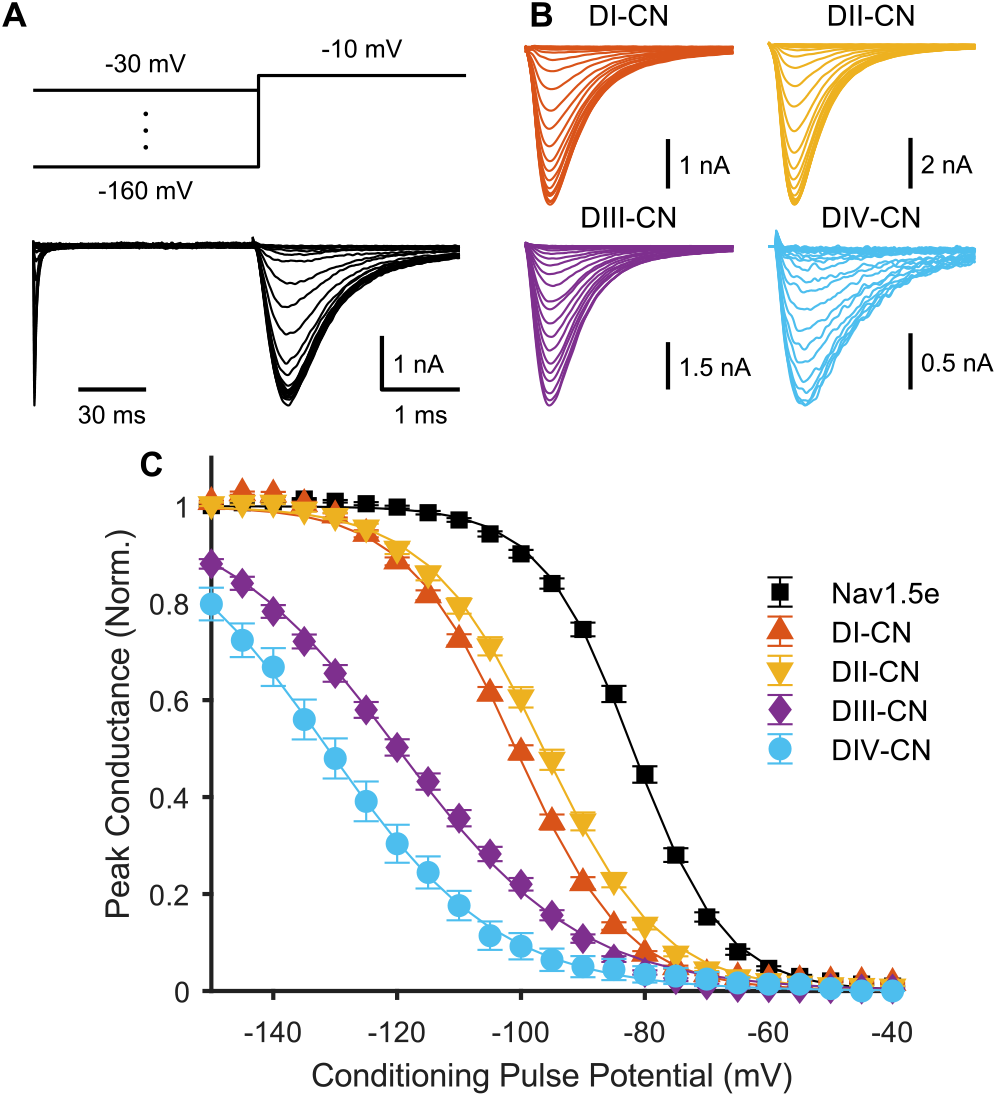
Impact of neutralizing voltage-sensing domains on channel inactivation. (**A**)Top: voltage step protocol used to assess steady-state inactivation (SSI). Conditioning pulses ranging from −160 to −30 mV were applied prior to a test pulse to −10 mV. Bottom: representative traces of ionic currents through WT Nav1.5e (cell 20170418c3) elicited by voltage protocol shown above. (**B**) Representative traces of currents through the four charge-neutralized mutants (DI-CN, cell 20180307c1; DII-CN, cell 20180316c1; DIII-CN, cell 20180712c3; DIV-CN, cell 20190318c3) following the test pulse to −10 mV. (**C**) Summary data corresponding to panel **A** and **B**, showing normalized peak current following the test pulse, as a function of the conditioning pulse voltage. Solid lines are fitted Boltzmann curves (Table S1).

### Voltage-Clamp Fluorometry Reveals That Domains III and IV Are Necessary for Inactivation

The charge neutralization experiments suggest that, at least for Nav1.5, both DIII and DIV may be involved directly in inactivation, since their neutralization led to both a dramatic hyperpolarization of SSI and a decreased sensitivity of inactivation to changes in membrane potential. Although a role for DIII in inactivation has been proposed previously (Armstrong, 2006; Armstrong and Hollingworth, 2018; Cha et al., 1999), charge neutralization data of Nav1.4 channels suggests that DIV alone is sufficient for inactivation (Capes et al., 2013). The argument favouring sufficiency is that neutralizing DIV in Nav1.4 leads to a shift in SSI that is so hyperpolarized, domains with intact voltage sensors must be in their deactivated state (Ahern et al., 2016; Capes et al., 2013). However, recent VCF data has shown that DIII activates at more hyperpolarized potentials in human Nav1.5 channels than in Nav1.4 channels (Chanda and Bezanilla, 2002; Hsu et al., 2017). To test whether this is also the case for the neonatal Nav1.5 used in this study, we quantified the intrinsic voltage-sensitivity of each domain in Xenopus oocytes using VCF. Fluorescent probes were conjugated to each domain, thereby engineering four domain-tagged VCF constructs: DI*, DII*, DIII* and DIV*. Steady-state fluorescence was then measured at voltage steps ranging from −150 mV (−180 mV for DIII*) to 50 mV.

In agreement with other VCF studies of Nav1.5 channels, the fluorescence-voltage (F-V) curve for DI* was more hyperpolarized than that for DII* (Fig. 4A, B) (Hsu et al., 2017; Varga et al., 2015). The voltage-dependence of the fluorescence signal of both DI* and DII* exhibited V_1/2_ values of −65.4 ± 2.3 mV (n = 13) and −44.1 ± 0.7 mV (n = 17) (see also, Table S1), respectively, which were approximately 10 mV more depolarized than V1/2 values reported for adult Nav1.5 channels (Hsu et al., 2017; Varga et al., 2015). This difference is in keeping with the more depolarized threshold for activation of Nav1.5e channels compared to the adult Nav1.5 splice variant (Onkal et al., 2008). Interestingly, the F-V plot of DII* had a shallower slope (k = 22.2 ± 0.6, n = 17) compared to DI* (k = 12.3 ± 1.2, n = 13), indicating that DII has a lower sensitivity to changes in membrane potential (Fig. 4A, B).

**Figure 4.**
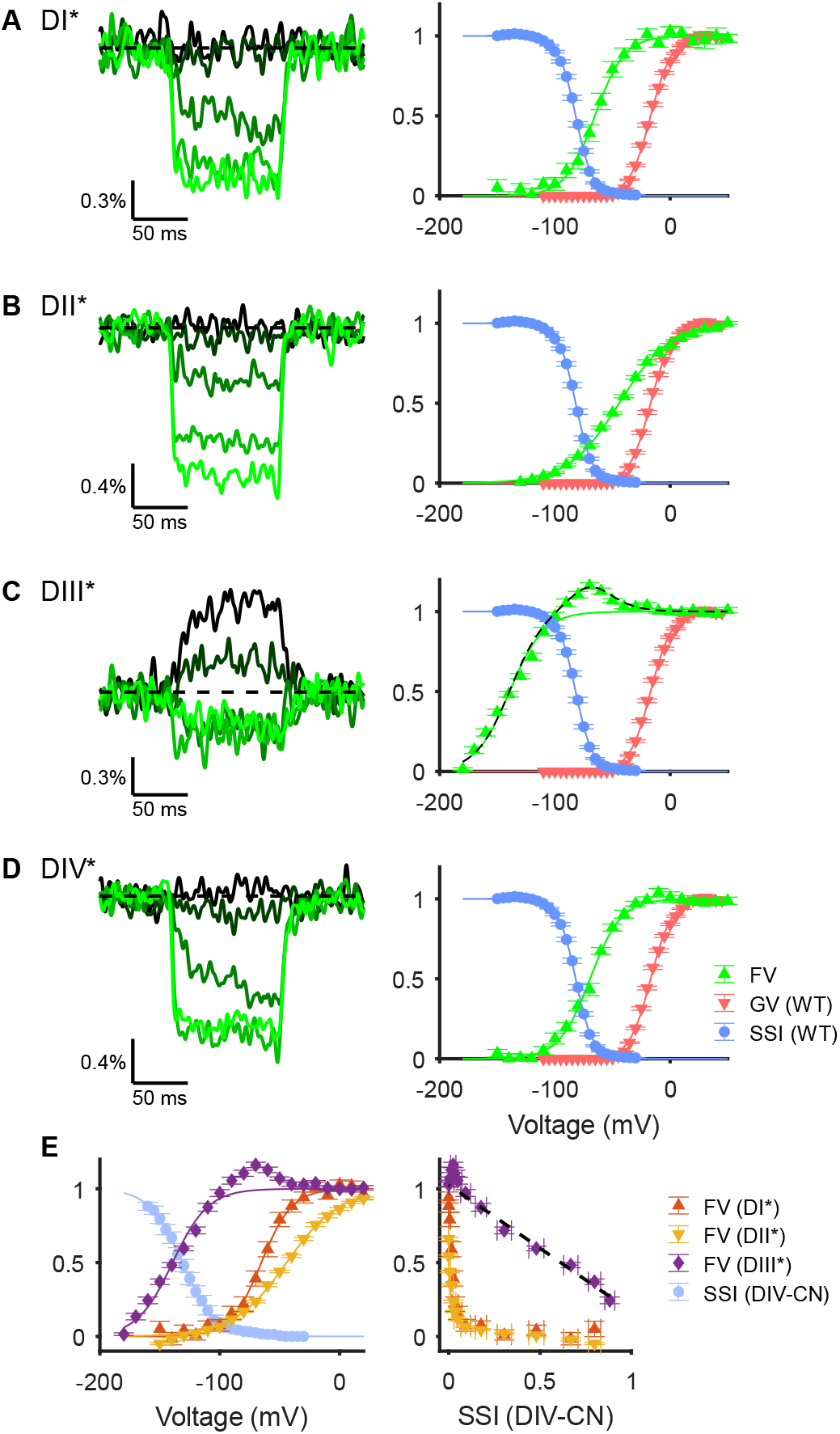
Voltage-dependence of fluorescence from tagged voltage-sensing domains. (**A**) Left: Representative fluorescence signals from a DI-tagged VCF construct (cell 20180510c2), recorded in Xenopus oocytes, in response to voltage steps of −140, −100, −60, −20 and +20 mV, shown by colour from darkest to lightest hue. Right: voltage-dependence of normalized fluorescence change from baseline (green circles). Solid green line is a fitted Boltzmann curve (Table S1). Overlaid is the SSI (blue triangles) and GV curve (red inverted triangles) of WT Nav1.5e channels (Fig. 1 and Fig. 3). (**B**) Same as panel A, but for DII-tagged VCF constructs. (Representative traces from cell 20180512c1). (**C**) Left: Representative traces for DIII-tagged VCF constructs (cell 20180518c4) in response to voltage steps of −180, −140, −100, −60, and −20 mV. Right: The dashed black line is the sum of the solid green line plus the derivative of the solid green line from panel **D**. (**D**) Same as panel **A**, but for DIV-tagged VCF constructs. (Representative traces from cell 20180514c5). (**E**) Left: Light blue circles represent the SSI curve from DIV-CN mutants (Fig. 3) overlaid with the fluorescence-voltage relationship of VCF constructs DI*-DIII* (panels **A**-**C**). Right: fluorescence change from baseline plotted against the SSI curve of DIV-CN. Line of best fit for DIII* versus DIV-CN SSI is shown as dashes black line (R^2^ = 0.97).

The normalized F-V relationship for DIII* was significantly more hyperpolarized than DI* and DII* with a V_1/2_ value of −137.4 ± 1.1 mV (n = 27) and a slope factor of 15.9 ± 0.6 (Fig. 4C). Interestingly, the fluorescence signal was biphasic, reaching a maximum at about −50 mV and declining in intensity at more depolarized potentials (Fig. 4C). This finding suggests that the voltage sensor of DIII may exhibit two distinct movements, analogous to the dynamics of the voltage sensor reported for Shaker K+-channels (Cha and Bezanilla, 1997) and DIII of Nav1.4 channels (Cha et al., 1999).

The F-V relationship observed for DIV* fluorescence occurred over a similar voltage range as SSI in WT channels (Fig. 4D) in keeping with the role of this domain in inactivation. The V_1/2_ value was −69.5 ± 1.3 mV (n=15) and the slope factor was 13.2 ± 0.5 mV. Interestingly, at the potentials at which we observed DIV movement, the F-V relationship of DIII* deviates distinctly from a Boltzmann function. We observed that the fluorescence change of DIII was well fit by a Boltzmann function plus the derivative of the DIV* Boltzmann fit (Fig. 4C). This observation might suggest an interaction between DIII and DIV movement, or between DIII and the binding of the inactivation motif.

Finally, the voltage dependence of the F-V plot for DIII* was strongly correlated with measurements of steady-state inactivation in DIV-CN mutants (Fig. 4E, left). In fact, plotting the SSI of DIV-CN mutants versus the fluorescence signal of DIII* at each membrane potential displayed a strong linear correlation (Fig. 4E, right). In contrast, the F-V curves for DI* and DII* were too depolarized to be correlated to SSI in DIV-CN mutants (Fig. 4E). This finding indicates that in the absence of DIV gating charges, DIII movement determines the voltage-dependence of SSI. Together with our previous results (Fig. 3C), this observation strongly suggests that in Nav1.5, both DIII and DIV are intrinsically necessary for channel inactivation, whereas DI and DII are not.

### Order of Voltage Sensor Movement Determines Closed-State Inactivation

The identified role of DIII in inactivation explains why DIII-CN affects SSI similarly to DIV-CN (Fig. 3C). We were thus left with two unexplained observations: why does neutralizing DI and DII hyperpolarize SSI (Fig. 3C) if they are not involved in inactivation (Fig. 4), and why does neutralizing DIV affect activation (Fig. 1C) if DIV movement is not necessary for pore opening (Fig. 2)? Since these effects did not occur in Nav1.4 channels (Capes et al., 2013), to understand our observations, we compared the outcomes of our reparametrized Nav1.5 model (Table S2) to the Capes *et al*. Nav1.4 model (Capes et al., 2013). We simulated DII-CN by removing the leftmost column of the model, consisting of the states C0, C4, and I4 (Fig. 5A). This is equivalent to assuming that the first step in the activation process is already complete; that is, the relevant domain has been biased towards its “active” conformation. Removal of states C0, C4, and I4 caused a leftward shift in the SSI curve without affecting activation in the Nav1.5 model (Fig. 5B). This shift in SSI was due to positive cooperativity between gating transitions that lead to channel activation and movement of the DIV voltage sensor (x_α_/x_β_>1, see Table S2). Notably, such cooperativity between voltage sensors has been reported in past VCF studies (Campos et al., 2007; Chanda et al., 2004). As a result of this cooperativity, neutralizing DI and DII increases the probability of inactivation directly from closed states, i.e. closed-state inactivation (CSI), even though they are not themselves necessary for inactivation.

**Figure 5.**
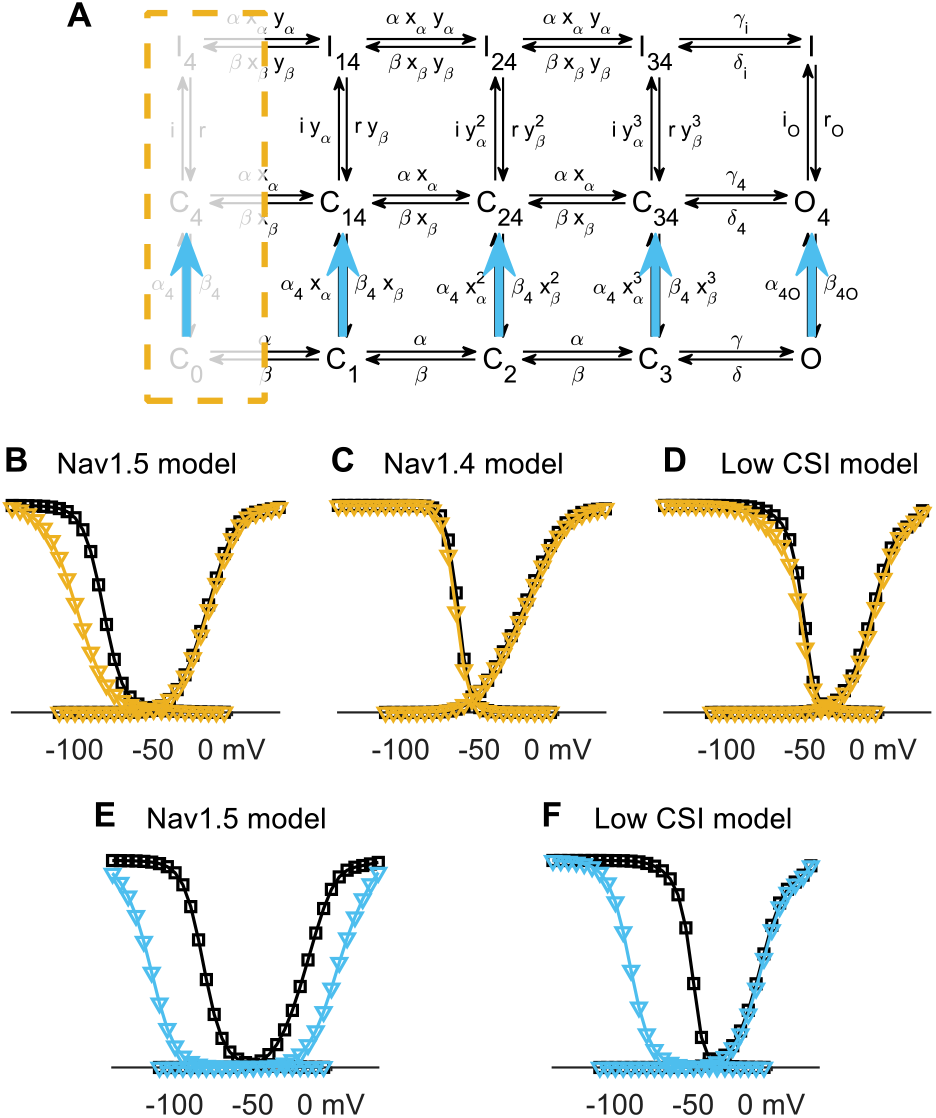
A kinetic model explains differences between Nav1.5 and Nav1.4. (**A**)Schematic of two separate modifications to the gating model. (Yellow) The leftmost column of states of the model has been removed, which simulates the charge neutralization of a non-specific domain whose movement is necessary for pore opening. (Blue) The rate α_4_ and α_4O_ have been increased 20-fold and β_4_ and β_4O_ have been decreased 20-fold, which approximates the removal of the bottom row, simulating the neutralization of DIV. (**B**) The peak activation (GV) curve (right) and steady-state inactivation (SSI) curve (left) produced by the model prior to (black) and following (yellow) the removal of the leftmost column of states. (**C**) Same as panel **B** but simulated with the Nav1.4 parameters from Capes *et al*. (2013). (**D**) Same as panel **B** but simulated with the modified Nav1.5 model that has a lower propensity for CSI, i.e. the “low CSI model” (see Methods). (**E**) GV and SSI curves produced by the model prior to (black) and following (blue) the approximate removal of the bottom row of states. (**F**) Same as panel **E** but simulated with the low CSI gating model (as in panel **D**).

Repeating our numerical experiment with the Nav1.4 parameters estimated by Capes et al. (Capes et al., 2013) revealed that the Nav1.4 model predicted their experimental observations, i.e. SSI was not affected (Fig. 5C). This was not because of a lack of interdomain coupling. In fact, the Nav1.4 model has similar levels of interdomain coupling as the Nav1.5 model (Nav1.4: x_α_/x_β_ =8.33, Capes et al., 2013; Nav1.5: x_α_/x_β_ =8.01, Table S2). However, we noted that in the Nav1.4 model, DIV moved exclusively following DI-III, whereas this was not true in the Nav1.5 model. Since DIV movement is necessary for inactivation, if DIV moves exclusively following pore opening, then CSI is not possible. To confirm that this difference was sufficient to explain the outcomes of neutralizing DI and DII in Nav1.4 and Nav1.5 channels, we modified the Nav1.5 parameters to force DIV to move only after pore opening (see Methods). Indeed, simulating DII-CN in this modified “low CSI” Nav1.5 model did not appreciably alter gating (Fig. 5D), consistent with experiments on Nav1.4. Altogether, these results suggest that Nav1.5 channels have an intrinsically higher propensity for CSI compared to Nav1.4 channels, which in turn modulates the functional consequences of neutralizing DI and DII.

We next simulated the effects of DIV-CN. Removing the entire bottom row of the model led to unrealistic conductance profiles (data not shown). We therefore elected to simulate DIV-CN by drastically biasing the transition probability from the bottom row to the middle row, by increasing the rates α_4_ and α_4O_ 20-fold and decreasing the rates β4 and β_4O_ 20-fold (Fig. 5A). In the Nav1.5e model, simulating DIV-CN led to a hyperpolarization of the SSI curve and depolarization of the GV curve (Fig. 5E), consistent with our experimental observations from Nav1.5e DIV-CN mutants (Fig. 1C and Fig. 3C). In the modified “low CSI” model, the same manipulation hyperpolarized SSI, with no effect on channel activation (Fig. 5F), again consistent with the effects of neutralizing DIV in Nav1.4 (Capes et al., 2013). These observations can be explained as follows. In the Nav1.5e (high CSI) model, the voltage-dependence of activation is set by a competition between pore opening and (closed-state) inactivation. When DIV is neutralized, inactivation is faster, shifting the balance towards CSI and thus requiring more depolarized voltages for pore opening to outcompete inactivation. In the low CSI gating model, activation is much faster than inactivation at all voltages, such that the voltage-dependence of activation represents the steady-state probability of pore opening, and thus is largely unaffected by a faster inactivation rate. In summary, both of the aforementioned differences in Nav1.4 and Nav1.5 CN mutants can be explained by differences in their intrinsic propensities for CSI.

### Auxiliary β-Subunits Selectively Target Closed-State Inactivation

The above results demonstrate that the functional contributions of each voltage sensor to channel gating depends both on the intrinsic coupling of the voltage sensor to the activation and/or inactivation processes *and* the sequential order of voltage sensor movements. In particular, we found that, with the exception of DIII, the voltage sensors in Nav1.4 and Nav1.5 are intrinsically coupled to essentially the same gating processes; however, in Nav1.5 a disposition for early DIV movement, and thus CSI, alters the consequences of voltage sensor neutralization. To extend these results beyond voltage sensor neutralization, we used the high and low CSI gating models developed in the previous section to analyze a range of more general gating perturbations, and found that the high and low CSI models responded in fundamentally different ways (Fig. S4). For example, stabilizing the resting state of DIV in the low CSI model did not alter activation or inactivation, whereas in the high CSI model the same perturbation depolarized SSI (Fig. S4). This was intriguing, because β1 and β3 subunits are thought to stabilize the resting state of DIV (Zhu et al., 2017). Our results thus predict that β1 and β3 subunits should differentially regulate high and low CSI channels, not because β1 and β3 interact with the voltage sensors differently, but simply because their CSI levels are different. This would suggest an unappreciated, physiological role for the sequence of voltage sensor movements in that it determines CSI levels. To test this, we sought to investigate the effects of β1 and β3 on Nav channels with different levels of CSI.

We first collected data from three different Nav channels: namely skeletal muscle Nav1.4 channels, cardiac Nav1.5 channels, and neuronal Nav1.6 channels (Fig. 6A, B). To experimentally quantify their propensities for CSI, we adapted the method proposed by Armstrong (Armstrong, 2006); briefly, the total conductance observed during a conditioning pulse was compared to the fraction of available channels during a subsequent test pulse (Fig. 6C, see also Methods). This analysis estimates the amount of open-state inactivation (OSI) occurring during the conditioning pulse, with CSI being defined as the fraction of SSI that cannot be accounted for by open state inactivation (OSI) (Fig. 6D). The estimated fraction of SSI occurring from closed states for Nav1.5e peaked at 100% and remained high for a large range of voltages (Fig. 6D, E). This was consistent with the Nav1.5e gating model where CSI can be determined exactly by adding up all the inactivation that occurs from closed states. To quantitatively compare CSI levels between isoforms, we fit the CSI curve with the difference of two Boltzmann curves; the integral of this function was defined to be the total amount of CSI. This analysis revealed that the three isoforms exhibited distinct levels of CSI (Fig. 6F), with Nav1.5e displaying the most, followed by Nav1.4 and finally Nav1.6 (Fig. 6F).

**Figure 6.**
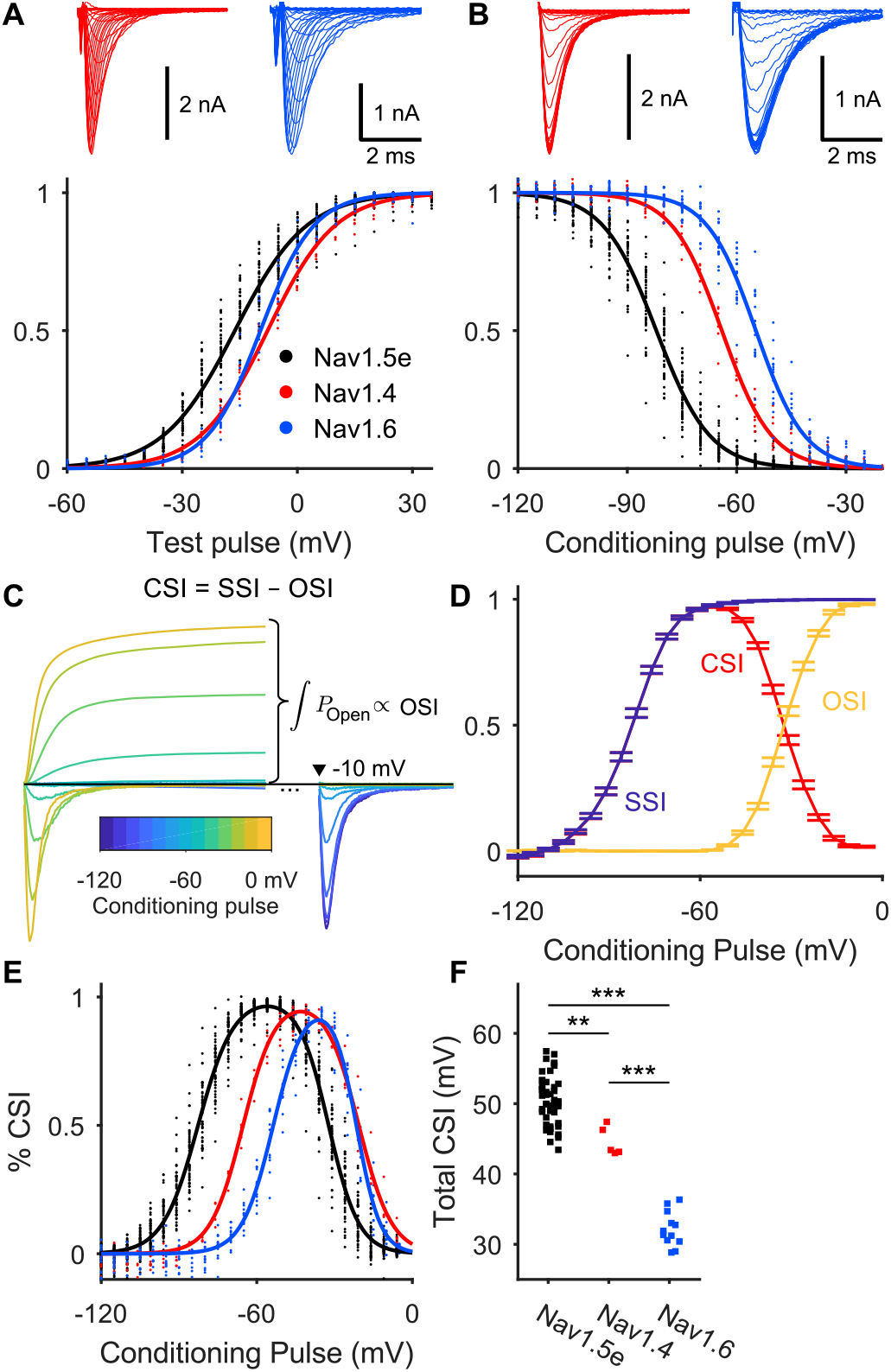
Comparison of closed-state inactivation (CSI) in different Nav isoforms. (**A**) Top: representative traces of Nav1.4 (red; cell 20181205c3) and Nav1.6 (blue; cell 20170419c4) activation. Bottom: peak conductance (GV) plots of Nav1.5e (black), Nav1.4 (red) and Nav1.6 (blue).(**B**) Top: representative traces of Nav1.4 (red) and Nav1.6 (blue) inactivation. Bottom: steady-state inactivation (SSI) plots. (**C**) Schematic showing how open-state inactivation (OSI) is calculated. The conductance of the channel during each conditioning pulse is summed to get the total probability of channel opening prior to the test pulse, which is proportional to OSI (see Methods for details). The difference between SSI and OSI reflects the fraction of channels that inactivated without conducting (i.e. CSI). Error bars represent mean ± S.E. (**D**) The fraction of Nav1.5e SSI (blue) due to OSI (yellow) and CSI (red) at each conditioning pulse potential. (**E**) Probability of CSI at different conditioning pulse potentials for each Nav isoform: Nav1.5e (black), Nav1.4 (red), and Nav1.6 (blue). The dots represent individual samples and the solid lines are the difference between two Boltzmann functions fitted to the data. (**F**) Area under the fitted CSI curves for each replicate. (**) indicates a significant difference between the group means with *p* < 10^−3^; (***) indicates *p* < 10^−6^.

If β1 and β3 regulate gating by stabilizing the resting state of DIV (Zhu et al., 2017), we would expect each of these isoforms to be differentially regulated, due to their distinct levels of CSI (see above). Coexpression of each isoform with either β1 or β3 confirmed this hypothesis. β1 and β3 led to depolarizing shifts in SSI of Nav1.5e, with no effects on channel activation (Fig. 7A). Consequently, the amount of total CSI was significantly reduced by both β-subunits (Fig. 7B). Nav1.4, which in our hands exhibited moderate levels of CSI (Fig. 6F), displayed moderate depolarizing shifts in SSI when coexpressed with β1 or β3 (Fig. 7C). CSI was reduced (Fig. 7D), but to a lesser degree than for Nav1.5e. Finally, activation and inactivation of Nav1.6 was unaffected by β-subunit coexpression (Fig. 7E), and the amount of total CSI was not altered (Fig. 7F). In summary, β1 and β3 depolarized SSI in each channel to a degree commensurate with the level of CSI displayed by the α-subunit alone (Fig. 6 and Fig. 7A-F), validating the regulatory mechanism proposed above.

**Figure 7.**
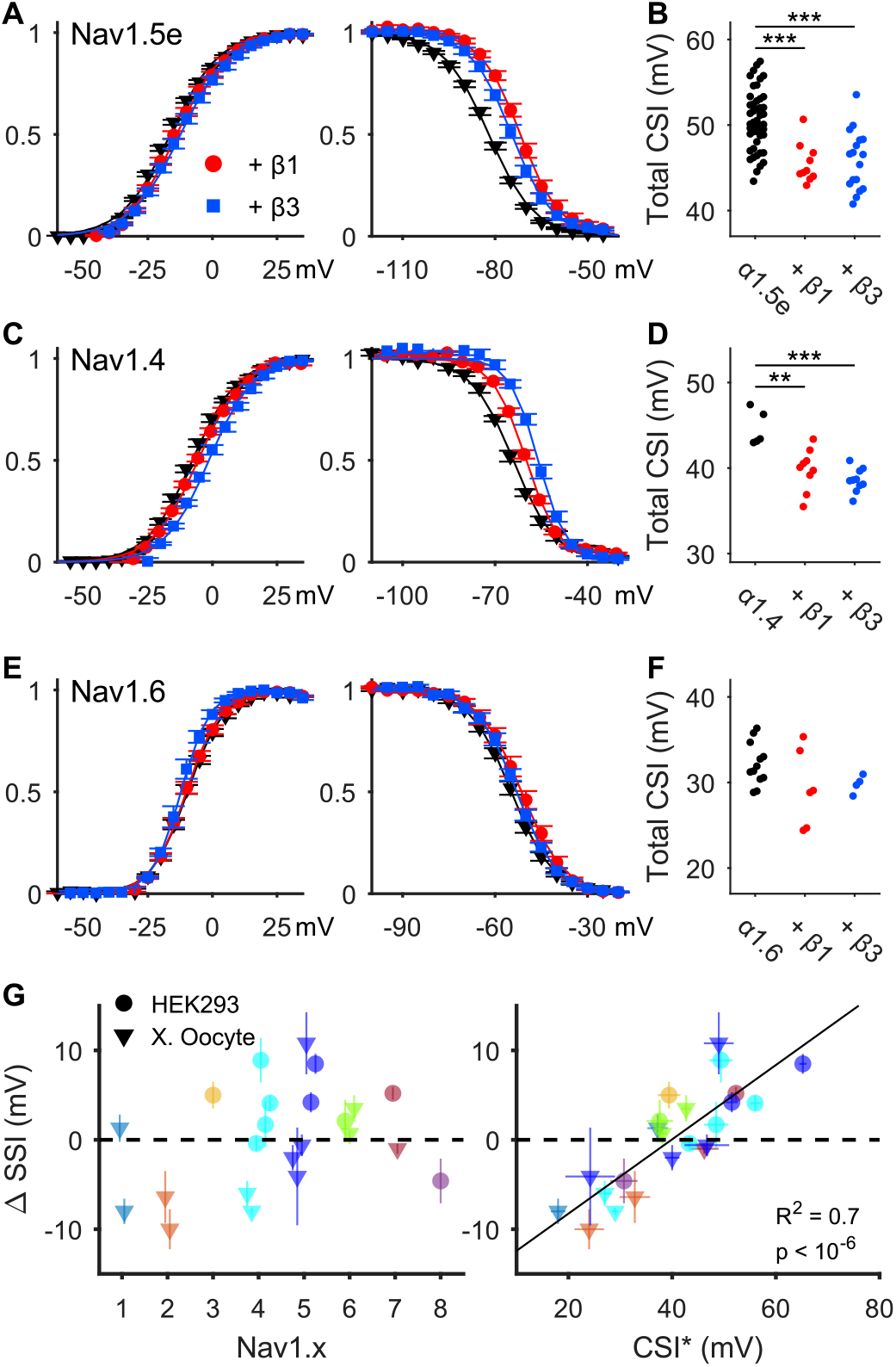
Comparison of β-subunit regulation on various Nav isoforms. (**A**)Left: peak activation as a function of voltage (G/V) for Nav1.5e alone (black, inverted triangles), and coexpressed with β1 (red, circles) and β3 (blue, squares). Right: steady-state inactivation (SSI).(**B**) Total CSI, as in Fig. 6F, for Nav1.5e alone and when coexpressed with β1 and β3. Each dot represents a sample. (***) indicates that *p* < 10^−4^. **(C-F)** Same as panel A and B, but for Nav1.4 and Nav1.6. (**) indicates p < 10^−2^. (**G**) Left: Change in the difference between reported V_1/2_ values for activation and steady-state inactivation (ΔSSI) following coexpression of β1 with various Nav isoforms (Nav1.1-Nav1.8). Data taken from past publications (Table S3). Circles represent data from mammalian cells, while triangles represent data from Xenopus oocytes. Lines represents standard error. Right: ΔSSI is plotted against CSI* measurement from the α-subunit alone. Nav isoform is colour-coded as in the left plot. R^2^ and p value for the line of best fit is reported in the bottom right.

It is well known that the effects of β-subunits on channel gating varies substantially between studies. We asked whether this variability could be explained by differences in CSI. To do this, we compared the reported effects of β1 on Nav1.1-Nav1.8 (Table S3). (We did not find sufficiently many studies to perform the same analysis for β3.) This cross-study comparison did not reveal strong evidence for β1 altering the voltage-dependence of activation (Table S3). However, the voltage-dependence of SSI exhibited large shifts following β1 coexpression (Fig. 7G, left). These shifts were extremely variable across studies, even between studies on the same α-subunit (Fig. 7G, left), suggesting that the varied effects of β1 are not due to isoform differences. Since we could not calculate CSI without access to the current recordings, we defined a “naïve” measure of CSI for each report as the difference between the V_1/2_ of activation and SSI, which we denote CSI* (Fig. 7G, right). CSI* of the α-subunits alone varied significantly across studies (Fig. 7G, right), but was nevertheless linearly correlated with the shifts in SSI following coexpression with β1 (R^2^=0.7). These observations indicate that the regulation of all Nav channel isoforms by β1 is determined by CSI. Altogether, these results suggest that the seemingly paradoxical reports of β1 regulation in the literature are, in fact, consistent with a singular mechanism of action.

## Discussion

The present study advances our understanding of voltage-gated sodium (Nav) channels in several important and interrelated ways. First, we show that the functional contributions of the various voltage sensing domains to channel gating depend on the channel’ s propensity for closed-state inactivation (CSI), a gating property which reflects the movement order of the voltage sensors. Second, this study reveals that Nav channels do not follow a single, prototypical gating sequence. Instead, our experiments demonstrate that the gating of Nav channels is better explained as a continuum of gating behaviours defined by each channel’ s propensity for CSI. Third and finally, we show that CSI is selectively targeted by β1 and β3 auxiliary subunits to exert their allosteric effect on skeletal muscle, cardiac, and neuronal Nav channels. In fact, we provide strong evidence that this mechanism extends to every Nav channel isoform, suggesting a novel form of channel regulation. In sum, we propose that the weak to strong modulation of all Nav channel isoforms by β1 and β3 is not a direct result of structural differences, per se, but is rather a natural consequence of variability in the intrinsic dynamics of channel gating.

### Differing contributions of voltage sensing domains to Nav channel activation and inactivation

Previous charge neutralization experiments have reported disparate changes to the voltage-dependence of activation when comparing skeletal muscle (Capes et al., 2013; Chahine et al., 1994), neuronal (Kontis and Goldin, 1997), and cardiac (Chen et al., 1996) Nav channels. Here, we show that changes to the voltage-dependence of activation is an unreliable marker for a domain’ s role in activation, whereas the latency to peak current is more informative (Fig. 2). Using this measure, we found that in the neonatal sodium channel, Nav1.5e, DI movement is likely rate limiting for pore opening and DII movement contributes significantly, but to a lesser degree. Although DIII movement is thought to be necessary for channel activation in Nav1.4 skeletal muscle channels (Chanda and Bezanilla, 2002), neutralizing DIII in Nav1.5 did not alter channel activation (Figs. 1 & 2). This is likely because DIII is already in its activated position in wildtype Nav1.5 channels at the voltage range used in our protocols (Fig. 4) (Hsu et al., 2017; Zhu et al., 2017). Consequently, neutralizing the domain’ s voltage sensor would not be expected to significantly alter the overall activation process. Finally, although DIV-CN Nav1.5 mutants exhibit an altered voltage-dependence of activation, the observed effects are more consistent with an increased inactivation rate (Fig. 2F & Fig. S4). Considering this, we did not find evidence that DIV movement is necessary for activation, consistent with previous experiments on Nav1.4 channels that were mutated to prevent inactivation (Goldschen-Ohm et al., 2013). In summary, our results suggest that DI and DII movements determine the rate of activation, whereas DIII, at least in Nav1.5, likely plays little role in activation at physiological membrane potentials.

Charge-neutralization experiments on Nav1.4 have suggested that DIV is uniquely sufficient for inactivation (Capes et al., 2013), whereas recent VCF experiments on Nav1.5 have strongly implicated both DIII and DIV in inactivation (Hsu et al., 2017). Here, we found that neutralizing either DIII or DIV led to a similar hyperpolarizing shift and flattening of the SSI curve. Furthermore, we observed that SSI in DIV-CN Nav1.5 mutants is strongly correlated with DIII movement (Fig. 4E). Together, this suggests that the Nav1.5 inactivation is intrinsically coupled to DIII movement. Notably, DIV-CN Nav1.4 mutants inactivate at voltages too hyperpolarized to be caused by the movement of DIII (Capes et al., 2013; Chanda and Bezanilla, 2002), and DIII-CN Nav1.4 mutants did not display significantly altered SSI (Capes et al., 2013). Accordingly, we conclude that DIII is coupled to inactivation differently in Nav1.4 and Nav1.5 channels, which could explain the divergent views surrounding the role of DIII in inactivation (Ahern et al., 2016; Armstrong and Hollingworth, 2018). Finally, although we found that in Nav1.5 neutralizing DI and DII significantly affected inactivation, these observations could be explained by interdomain coupling with DIV. Nevertheless, our observations suggest that DI and DII movements can functionally contribute to channel inactivation, even if they are not necessary for inactivation, per se. Overall, our results highlight unappreciated variability in the molecular basis of channel gating across Nav channel isoforms.

### Closed-state inactivation modulates the contributions of voltage sensing domains

It is well established that Nav channels can inactivate from closed states (Aldrich et al., 1983; Armstrong, 2006; Bean, 1981; Lawrence et al., 1991; Vandenberg and Horn, 1984), which is thought to occur when DIII and DIV (possibly just DIV in Nav1.4) move prior to channel activation (Armstrong, 2006). Nevertheless, the prevailing idea is that DIV movement is substantially slower than the other domains across all voltages (Armstrong, 2006; Bosmans et al., 2008; Capes et al., 2013; Chanda and Bezanilla, 2002), implying that inactivation occurs predominantly from the open state. Interestingly, almost all evidence for this delayed DIV movement is from Nav1.4 channels (Capes et al., 2013; Chanda and Bezanilla, 2002; Goldschen-Ohm et al., 2013). In contrast to Nav1.4, however, we observed a high propensity for CSI in wildtype Nav1.5 channels (Fig. 6D), indicating a gating sequence that is preferential to early DIV movement. We showed that such a difference in gating sequence is sufficient to explain the contrasting outcomes of voltage sensor neutralization between Nav1.4 and Nav1.5 channels. Furthermore, we found that this apparent dichotomy of low and high CSI gating in fact defines a *continuum* of gating behaviours along which all Nav channel isoforms lie (Fig. 7 & 8). This study thus suggests that the gating sequence is a variable parameter which modulates the contributions of each voltage sensor to activation and inactivation across Nav channels. More generally, our modelling predicts that CSI should modulate the effects of any gating modifiers which target the movements of these voltage sensing domains. Indeed, this mechanism succeeded in predicting the effects of auxiliary β subunits (discussed below), highlighting the unappreciated importance of CSI in the biological functionality of all Nav channels.

### Intrinsic gating properties of Nav channels dictate auxiliary subunit regulation

The auxiliary subunits, β1 and β3, have surprisingly varied effects on Nav channel gating, even within a single isoform (Fig. 7G, Table S3). Consequently, this variability is likely not due to differences in channel structure (Jiang et al., 2020; Shen et al., 2019). Rather, the observations made here motivate an alternative explanation: β1 and β3 specifically target CSI such that their effects on channel gating are determined by the variable dynamics of the pore-forming subunit. Theoretically, we found that if β1 and β3 stabilize the resting state of DIV, as demonstrated in Nav1.5 using VCF (Zhu et al., 2017), then their effects should depend on the channel’ s propensity for CSI (Fig. S4). This prediction was confirmed by our experiments comparing Nav1.4, Nav1.5, and Nav1.6 channels (Fig. 7).

Intriguingly, the idea that the intrinsic gating properties of a channel predetermine the effects of allosteric modifiers has also recently been suggested for AMPA receptors (Dawe et al., 2019). Depending on the channel splice variant (flip or flop), gating modifiers, such as anions and auxiliary subunits, either exert an effect (flip) or no effect (flop), similar to what we observe for high and low CSI Nav channels. This switching in regulation was shown to be due to the mobility of the apo or resting state of the receptor (Dawe et al., 2019), suggesting a critical role for the intrinsic dynamics of the pore-forming subunits. These observations point to an over-arching principle of ion channel regulation which could be tested by exploring the potential role of the intrinsic gating to other ion channel families.

Notably, our predictions concerning the influence of CSI on Nav channel regulation extend to any gating modifiers which target the movements of specific voltage sensing domains. Interestingly, several studies on toxins that target voltage sensors – which include certain scorpion, sea anenome, and cone snail toxins (Ahern et al., 2016) – have described varied effects across Nav channel isoforms (Alami et al., 2003; Leipold et al., 2006; Oliveira et al., 2004). Although many of these observations have been ascribed to structural variation between the channels, it would be informative to assess the extent to which differences in CSI may also contribute. Since any compounds which target voltage sensor movements in Nav channels have potential as novel drugs for chronic pain, epilepsy, and heart disorders (Bosmans and Swartz, 2010; Cardoso and Lewis, 2019), identifying sources of CSI variability *in vivo* and understanding why CSI is so poorly controlled for *in vitro* (Fig. 7G) may have particular importance for novel drug design. Finally, it has been suggested that insights into channelopathies may be realized through the analysis of “homologous” mutations across Nav channels (Loussouarn et al., 2016). Whether CSI alters the consequences of other mutations as profoundly as the voltage sensor neutralizing mutations performed here is clearly of interest for future study.

## Methods

### Molecular biology

The mouse mH1 pcDNA3.1(+)-plasmid was obtained from Dr. T. Zimmer (Camacho et al., 2006). Exon 6a cDNA was amplified out of mouse brain homogenates, using the Access RT-PCR System (Promega), and then exchanged with exon 6b in mH1 using the Quikchange method of site-directed mutagenesis (SDM) (Braman et al., 1996), to generate the Nav1.5e plasmid. A similar RT-PCR approach was also used to clone out the mouse β1 and β3 cDNA and insert it into a pcDNA3.1(+) vector.

Charge-neutralizing mutations of Nav1.5e were engineered using single-primer reaction in parallel (SPRINP) SDM (Edelheit et al., 2009). Primers (Integrated DNA Technologies) were designed containing both the amino acid exchanges of interest and a silent restriction site. Following the PCR reaction, unmutated templates were digested using 0.4-0.8 U/μL of DpnI (New England Biolabs). The resulting PCR mixture was transformed into house-grown competent DH5α. Colonies were grown overnight on agar plates and picked for liquid culturing in 25 g/L Lysogeny Broth (Fisher Bioreagents). Plasmids were harvested using QIAprep Spin Miniprep Kits (Qiagen). Mutations were screened via restriction digest and gel electrophoresis, using the silent restriction sites initially designed into the SDM primers. Plasmid sequences were verified with Sanger sequencing done by the Innovation Centre of McGill University and Genome Quebec, using Sequencher 4.8 and CLC Sequence Viewer 8.0.

Nav1.5e DIV-CN constructs were modified further by conjugating the Nav sequence to an engineered GFP fluorophore called Mystik (Mys) via a P2A linker. The P2A allowed for the stoichiometric 1:1 translation of the Mystik and Nav genes, so that cells showcasing the strongest fluorescence were the most likely to express the Nav channel in higher abundance (Ahier and Jarriault, 2014). The allowed us to overcome the poor expression of Nav1.4 alone. Primers (Integrated DNA Technologies) with overhangs containing fragments of the Nav plasmid’ s 5’ untranslated region or part of the P2A-Nav sequence were used to PCR amplify a double-stranded “megaprimer” containing (from 5’ to 3’): the 5’ UTR upstream of the Nav channel, the entire Mys gene, the P2A sequence, and the 5’ end of the Nav channel cDNA. This megaprimer was then isolated via gel electrophoresis, extracted into solution using MinElute Gel Extraction Kits (Qiagen), and inserted into the Nav channel plasmid by means of the Quikchange method of PCR discussed previously.

### Cell culture

Human embryonic kidney cells mutated to over-express the SV40 large T-antigen (HEK-293T cells) were used as the expression system for electrophysiological recordings. HEK-293T cultures were maintained in minimal essential media containing Glutamax (MEM-glutamax; Gibco), supplemented with 10% fetal bovine serum (FBS; Gibco), and kept at conditions of 37°C, 100% humidity, and 5% CO_2_ in a ThermoForma Series II Water-Jacketed CO_2_ Incubator (ThermoFisher). Main HEK-293T stocks were grown in T-25 Flasks (Corning) and twice a week passaged using Trypsin-EDTA solution (Gibco) into 35-mm Tissue-Culture Treated Culture Dishes (Corning) for transfection purposes. The calcium phosphate transfection protocol (Jordan et al., 1996) was used to transiently transfect HEK-293T cells at least 24 hours before recordings. Each 35-mm culture dish was loaded with 0.5 µg of the Nav channel plasmid, 0.2 µg of the transfection reporter Mys (unless already present, as in the Mys-P2A-Nav constructs), and, if present, 1.5 µg of β1 or β3. The cDNA mixture was dissolved in 560 mM of CaCl_2_ (unless otherwise indicated, all chemical reagents are from Sigma-Aldrich), and an equal volume of 2XBES solution (in mM: 50 BES, 280 NaCl, and 1.5 Na2HPO4) was added to induce precipitate formation. After roughly one minute, the DNA-calcium phosphate precipitates were added to a single culture dish. The dish was then returned to the incubator where cells had time to be transfected, typically over 6 to 9 hours. The reaction was quenched 6 to 9 hours later by rinsing with phosphate-buffered saline (PBS, in mM: 137 NaCl, 2.7 KCl, 10.1 Na_2_HPO_4_, and 2 NaH_2_PO_4_) containing 1 mM EDTA and rinsed with PBS containing 1 mM MgCl_2_ and 1 mM CaCl_2_. Transfected HEK-293T cells were left to recover overnight.

### Electrophysiology

At least two hours before the start of recordings, transfected cells were dissociated from their cultures using Accutase and then re-plated at a lower density. This step increased the yield of isolated cells, minimizing gap junctions that form between adjacent HEK-293T cells and optimizing the quality of the voltage-clamp conditions. Culture media was replaced with external solution containing (in mM): 155 NaCl, 4 KCl, 5 HEPES, 1 MgCl_2_, and 1.8 CaCl_2_, with pH adjusted to 7.3-7.4 using NaOH. Cells were patched with microelectrodes containing an internal solution that was optimized for voltage-clamp conditions, made up of (in mM): 115 CsCl, 5 HEPES, 5 Cs_4_-BAPTA, 1 MgCl_2_, 0.5 CaCl_2_, 10 Na_2_ATP, with pH adjusted to 7.3-7.4 using CsOH and sucrose added to keep the osmolality matching that of the external solution, between 295-300 mOsm. Borosilicate glass capillaries – with an inner diameter of 1.15 mm, an outer diameter of 1.65 mm, a length of 100 mm, and a 0.1 mm filament (King Precision Glass, Inc) – were pulled using a PP-830 vertical puller (Narishige), yielding microelectrodes with a pipette resistance of 1 to 5 MΩ. Microelectrode tips were then dipped into Bees-Wax Pure Natural (Integra Miltex) and subsequently fire-polished with an MF-900 Micro Forge (Narishige), to reduce noise and improve membrane seals. Dissociated cells were viewed using an Eclipse Ti-U Inverted Microscope (Nikon). Transfected cells were identified by their green fluorescence, excited by a DC4104 4-Channel LED Driver (Thorlabs). Positive pressure was applied orally to electrodes before being lowered to the bottom of the recording chamber using an MP-285 micromanipulator (Sutter Instrument). Once the electrode tip was positioned just above the surface of the cell, the release of the positive pressure was sufficient to form a glass-membrane seal of at least 1 GΩ. Pulses of negative pressure were then delivered to improve the strength of the seal and eventually break through, giving access to the whole-cell patch-clamp configuration. All recordings were done at room temperature.

Voltage commands were delivered through an AxoPatch 200B Amplifier (Axon Instruments). Capacitive transients from the pipette and from the cell were cancelled, cell capacitance and series resistance were monitored to avoid changes exceeding 30% over the course of the recording, and series resistance was compensated to the amplifier’ s maximum (98% on the machine, but in practice probably closer to 80%). Currents were acquired at 100 kHz, low-pass Bessel filtered at 5 kHz using a Model 900 Tunable Active Filter (Frequency Devices), and telegraphed via an Axon Digidata 1550 (Molecular Devices). All data was collected and saved digitally using pClamp 10.7 software (Axon Instruments).

### Voltage-clamp protocols

When cells were not being recorded, either between protocols or between sweeps within a protocol, they were clamped at a holding potential of −60 mV. To assess channel activation, cells were stepped down to the baseline potential for 300 ms, depolarized to a series of potentials between −110 and +70 mV in increments of 5 mV for 100 ms, then returned to the baseline potential for another 300 ms. This baseline potential was either −100 mV for Nav1.5e, or −130 mV in the case of the charge-neutralized mutants, to counter the latter’ s increased propensity for channel inactivation. To assess steady-state inactivation, cells were stepped down to the baseline potential for 300 ms, given a pre-pulse that varied between −160 mV and −5 mV in increments of 5 mV for 100 ms, pulsed directly to the test pulse potential of −10 mV for 50 ms, then returned to −100 mV for an additional 300 ms. Leak subtraction was performed on the raw data in a custom Igor Pro (Wavemetrics) program post-hoc.

### Voltage-clamp fluorometry

All experiments conducted on *Xenopus laevis* were done with the approval of and according to the guidelines established by the Animal Care Committee of the National Institutes of Natural Sciences, an umbrella institution of the National Institute for Physiological Sciences. We obtained the Nav1.5 VCF constructs used by Varga et al. (Varga et al., 2015) from Dr. Johnathon R. Silva and used site-directed mutagenesis, as described previously, to convert them to their respective Nav1.5e variants (with the exception of position 215, which was not mutated to leucine because it was used as a labelling cysteine). The four Nav genes, inserted on pMax vectors, could be linearized using Pacl (TOYOBO) and used as a template to generate cRNA using the mMESSAGE T7 RNA transcription kit (Thermo Fisher Scientific).

The oocytes of *Xenopus laevis* were surgically harvested from anaesthetized animals as described previously (Kume et al., 2018). Oocytes were separated ad defolliculated using 2 mg/ml collagenase treatment for 6.5 hours. Then oocytes were incubated overnight at 17°C in Ringer’ s solution containing (in mM): 88 NaCl, 1 KCl, 2.4 NaHCO_3_, 0.3 Ca(NO_3_)_2_, 0.41 CaCl_2_, 0.82 MgSO_4_, and 15 HEPES, with pH adjusted to 7.4 with NaOH and use of 0.1% penicillin-streptomycin. The remaining follicular layers were manually peeled. 25 ng of Nav channel cRNA was injected into the vegetal pole of oocytes using Nanoject II (Drummond). Treated oocytes were then returned to their 17°C incubator and left for 1 to 3 days to express protein.

Oocytes were labelled for 20 minutes using 10 μM of methanethiosulfonate-carboxytetrmethylrhodamine (MTS-TAMRA) dye on ice. Dye conjugation was done in depolarizing solution (containing, in mM, 110 KCl, 1.5 MgCl2, 0.8 CaCl2, 0.2 EDTA and 10 HEPES, pH adjusted to 7.1 with KOH) in an effort to expose and label S4 helices. Excess dye was removed by rinsing five times with fresh ND-96 solution (96 mM NaCl, 2 mM KCl, 1.8 mM CaCl2, 1 mM MgCl2, 5 mM HEPES, with pH adjusted to 7.4 wit NaOH). Oocytes in solution were kept on ice until use. Eventually, oocytes were transferred to the recording chamber filled with ND-96 solution, at room temperature, oriented in such a way that fluorescent recordings were done on the animal pole. Voltage clamp for macroscopic current recording was performed by using an amplifier (OC-725C, Warner Instruments). Borosilicate glass capillaries (World Precision Instruments) were used with a resistance of 0.1–0.3 MΩ when filled with 3 M KOAc and 10 mM KCl. The fluorescent recordings were performed with the fluorescence microscope (Olympus BX51WI) equipped with a water immersion objective lens (Olympus XLUMPLAN FL 20x/1.00). The light source was emitted by a xenon arc lamp (L2194-01, Hamamatsu Photonics) and passed through a band-pass excitation filter (520-550 nm). The intensity of the excitation light was decreased to prevent the fluorophore from bleaching by ND filters (Olympus U-25ND6 and U-25ND25). Emitted light was passed through a band-pass emission filter (580IF). The emission signals were detected by a photomultiplier (Hamamatsu Photonics H10722-110). The baseline signal was adjusted to 2 V. The detected current and fluorescent signal were acquired by a Digidata 1332 (Axon Instruments) and Clampex 10.3 software (Molecular Devices) at 100 kHz.

Oocytes were held at a baseline of −60 mV between sweeps and protocols. Once we could detect a visible change in fluorescence upon depolarization of the oocyte from −120 mV to +60 mV, we could run the following activation protocol. The oocyte was stepped down to −120 mV for about 300 ms, then hyperpolarized/depolarized to a series of potentials between −180 mV and +60 mV in increments of 10 mV for 100 ms, then stepped back to −120 mV for about 300 ms before returning to baseline. The protocol was run and averaged 10 times to improve the signal-to-noise ratio in the fluorescent output. Voltage-clamp fluorometry data was analyzed automatically using custom-made Igor Pro scripts.

### Literature search

Values for the effect of β1 coexpression on Nav gating were gathered from published studies found through searching keywords on Google Scholar and from papers cited in these studies. The search was restricted to data collected in either HEK293 cells or Xenopus oocytes. Following least-squares regression of ΔSSI against CSI*, one outlier was identified using Cook’ s distance and subsequently excluded from the analysis (Nav1.8; Xenopus oocytes; (Vijayaragavan et al., 2001)). All numerical values are reported in Table S3.

### Data analysis and statistics

For activation, E_Na_ was determined by fitting the linear part of the current-voltage curve to a line and extrapolating its x-intercept. Conductance was calculated as I/(V-E_Na_), then fit with a Boltzmann function

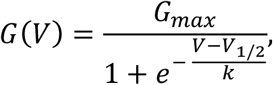

where G_max_ is the maximum conductance, V_1/2_ is the half-activation potential, and k is the slope factor of voltage sensitivity. For steady-state inactivation, peak currents following the test pulse were plotted against the corresponding pre-pulse potential; a Boltzmann function was then fit to this I-V curve. To calculate time to peak, the maximum current occurring at least 0.5 ms following the voltage step was identified (to avoid identifying capacitive transients at low voltages). The latency from the moment of voltage step to this peak was calculated.

To experimentally quantify the fraction of inactivation occurring from closed states (i.e. closed-state inactivation), we adapted the method proposed by Armstrong (Armstrong, 2006). Briefly, open-state inactivation was estimated during a conditioning voltage step (−100 ms long, ranging from −120 to −5 mV), and then compared to the fraction of available channels during a subsequent test pulse (−10 mV), thereby estimating the fraction of steady-state inactivation occurring through open states (Fig. 6C). Closed-state inactivation was then defined as the complementary fraction. Specifically, open-state inactivation (OSI) was estimated by the equation

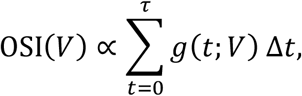

where *g*(*t*; *V*) is the experimentally observed channel conductance at time t for voltage step V. In the original method (Armstrong, 2006), OSI was scaled by assuming 100% of steady-state inactivation occurs from open states at 0 mV. However, our data displayed significantly more closed-state inactivation than was observed by Armstrong (Armstrong, 2006) and consequently OSI did not necessarily attain a maximum within the tested voltage range. We therefore modified this method by fitting a Boltzmann function, 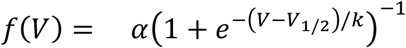, to the unscaled OSI versus voltage curve to obtain a scaling factor, i.e. α. This allowed open-state inactivation to attain its maximum outside of the observed voltage range.

For statistical analysis, Tukey’ s method was used following a one-way ANOVA, with a significance level of *p* < 0.01. All data was reported as mean ± SE. CSI* was calculated as the difference between group means; the standard error estimate of CSI* (for plotting purposes, see Fig. 7G) was thus defined in the sense of Welch’ s t-test, i.e. 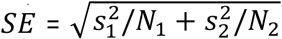.

### Computational modelling

For simulations of Nav gating, the kinetic model of Nav1.4 developed by Capes et al. (Capes et al., 2013) was reparametrized with a custom-made evolutionary fitting algorithm programed in MATLAB 2017b (The MathWorks, Inc.). The first generation of models were initialized with parameters drawn from a multivariate Gaussian centered at the Nav1.4 parameters with zero covariance and a coefficient of variation of 1. Each individual was assessed on z-scores calculated with respect to our experimental observations of the integral, peak amplitude, and time to peak of the activation and inactivation currents, as well as the recovery from inactivation, steady-state inactivation, and IV curves (Fig. S2). We ran the algorithm until the maximum fitness did not improve by at least 10% over 40 generations. When reparametrizing the model, we relaxed several assumptions made by the original authors. Specifically, we removed the constraint that the voltage-dependent charge of transitions reflecting movement of the same domain be the same and did not define any parameters with respect to others, no longer forcing microscopic reversibility. Nonetheless, the modelled conductances were well behaved during all simulations and our sensitivity analyses demonstrated a smoothness of model output with respect to local perturbations (Fig. S2E, F), suggesting that we did not overfit the model. The resulting parameter values are reported in Table S2.

To produce the low CSI gating model (Fig. 5 & Fig. S4), the parameters where adjusted to bias DIV to move following pore opening. This was done by decreasing the forward rate of DIV movement while the channel was closed (α_4_ was decreased 100 fold) and increasing the forward rate following pore opening (α_4O_ was increased 100 fold). Consequently, the probability of CSI was reduced.

All simulated protocols were identical to those applied experimentally. The modelled current was calculated as I_mem_ = (O + O_4_)(V − E_rev_), where E_rev_ is the average reversal potential calculated experimentally, O and O_4_ are the open states of the channel, and V is voltage. As for all protocols with applied step voltages, solutions to the model were computed using the exponential of the transition matrix.

## Data and materials availability

Summary data and all the code used in the generation of figures is publicly available at https://github.com/niklasbrake/Nav2020. Raw data is available upon request from the authors.

## Acknowledgments

We thank Dr. Mark Aurousseau who cloned Nav1.5e from brain extracts and Dr. Jonathan R. Silva for the gift of Nav1.5 voltage-clamp fluorometry cDNA constructs. We also thank members of the Bowie laboratory, for discussions and comments on the manuscript.

## Author contributions

Conceptualization: NB, ASM, and DB

Methodology: NB, TS, and YK

Software: NB

Validation: HS

Formal Analysis: NB, ASM, YY, HS, and TS

Investigation: NB, ASM, YY, and HS

Writing – Original Draft: NB and DB

Writing – Review & Editing: all authors

Visualization: NB

Supervision: AK and DB; Project Administration, DB Funding Acquisition: YK, AK, and DB

## Supplementary Figures & Tables

**Fig. S1.**
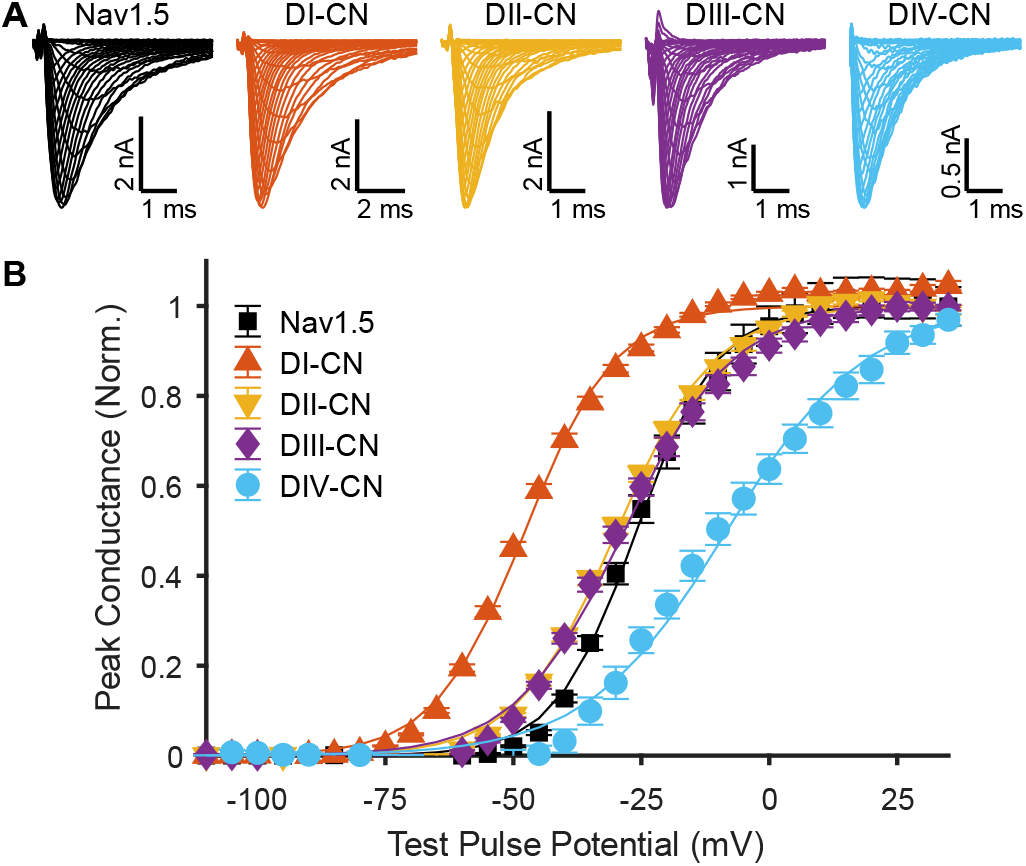
Changes in peak GV following domain neutralization in adult form of Nav1.5, mH1. (**A**) Representative traces of ionic currents corresponding to wild-type (WT) Nav1.5 (cell 20170311c3) and mutant channels (DI-CN, cell 20180301c6; DII-CN, cell 20180711c3; DIII-CN, cell 20180309c12; DIV-CN, cell 20181022c7) in response to depolarizing voltage steps ranging from −110 to 35 mV, following a holding potential of −130 mV (−100 mV for WT), recorded in HEK-293T cells. (**B**) Summary data corresponding to panel **A**, showing normalized peak current following the test pulse, as a function of the test pulse voltage.

**Fig. S2.**
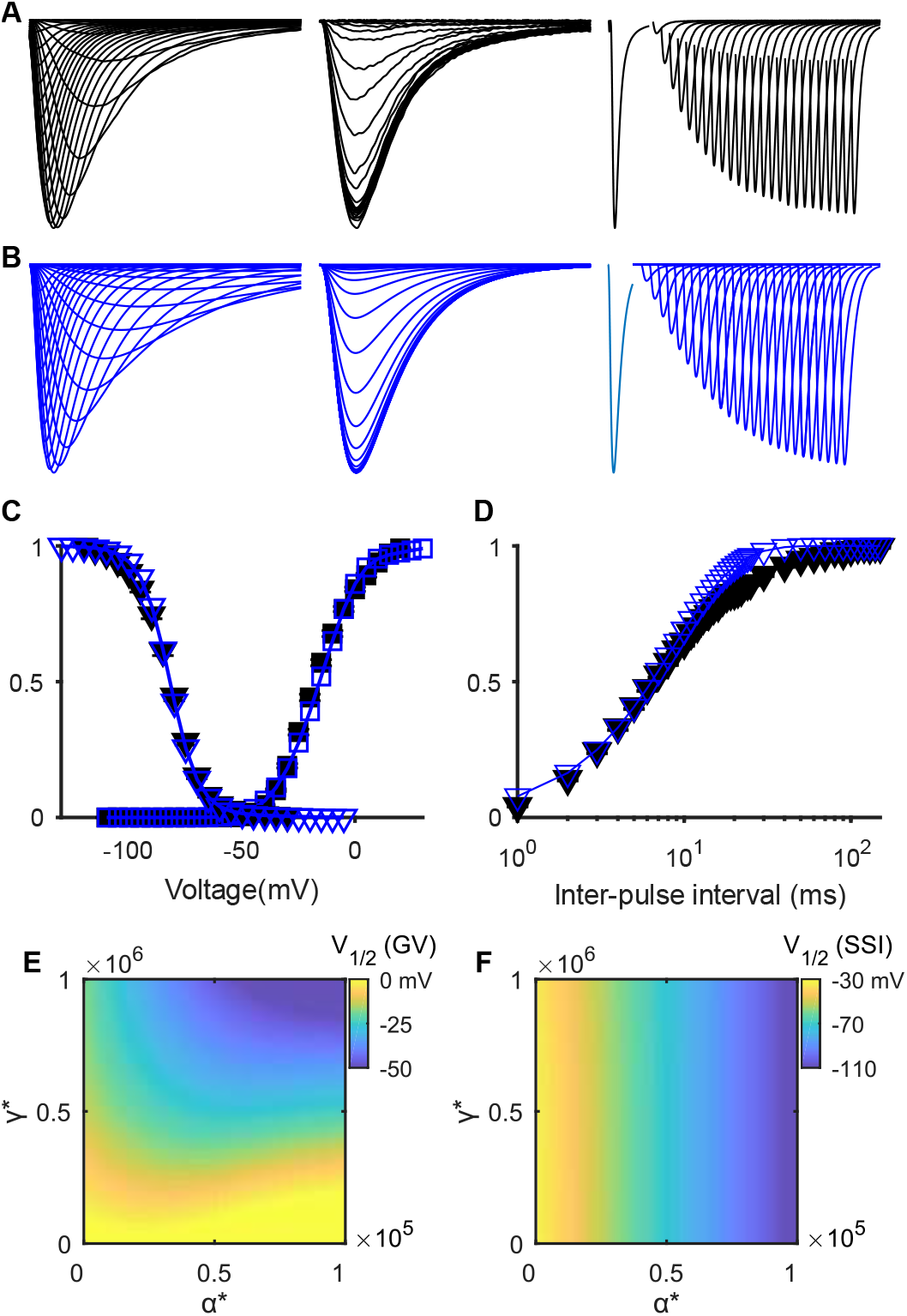
Comparison of reparametrized Nav1.5e model to data. (**A**) From left to right, averaged currents recorded from WT Nav1.5e channels during the activation protocol (as in Fig. 1), inactivation protocol (as in Fig. 3), and recovery from inactivation protocol (currents elicited by a test pulse of −10 mV, following a holding potential of −10 mV and a hyperpolarizing inter-pulse of −120 mV which lasted between 1-150 ms). (**B**) Modelled currents with the gating model in Fig. 2A and parameters from Supplementary Table S2. (**C**) Steady-state inactivation (inverted triangles) and GV (square) curves of WT Nav1.5e channel (black) and model (blue). (**D**) Fraction of recovered current as a function of inter-pulse interval for the WT channel (black) and model (blue). (**E**) Heat-map showing the sensitivity of the V_1/2_ of GV. At low values of α*, changes in γ* have little effect on the V_1/2_ of GV, and vice versa. However, as both are increased, the V_1/2_ of GV becomes more hyperpolarized. (**F**) Heat-map showing the sensitivity of the V_1/2_ of SSI to changes in α* and γ*. As α* increases, a hyperpolarizing shift in the V_1/2_ of SSI is observed, whereas changes in γ* produce almost no effects on SSI. Both heat-maps are color-coded based on the color-bars to the right of each panel.

**Fig. S3.**
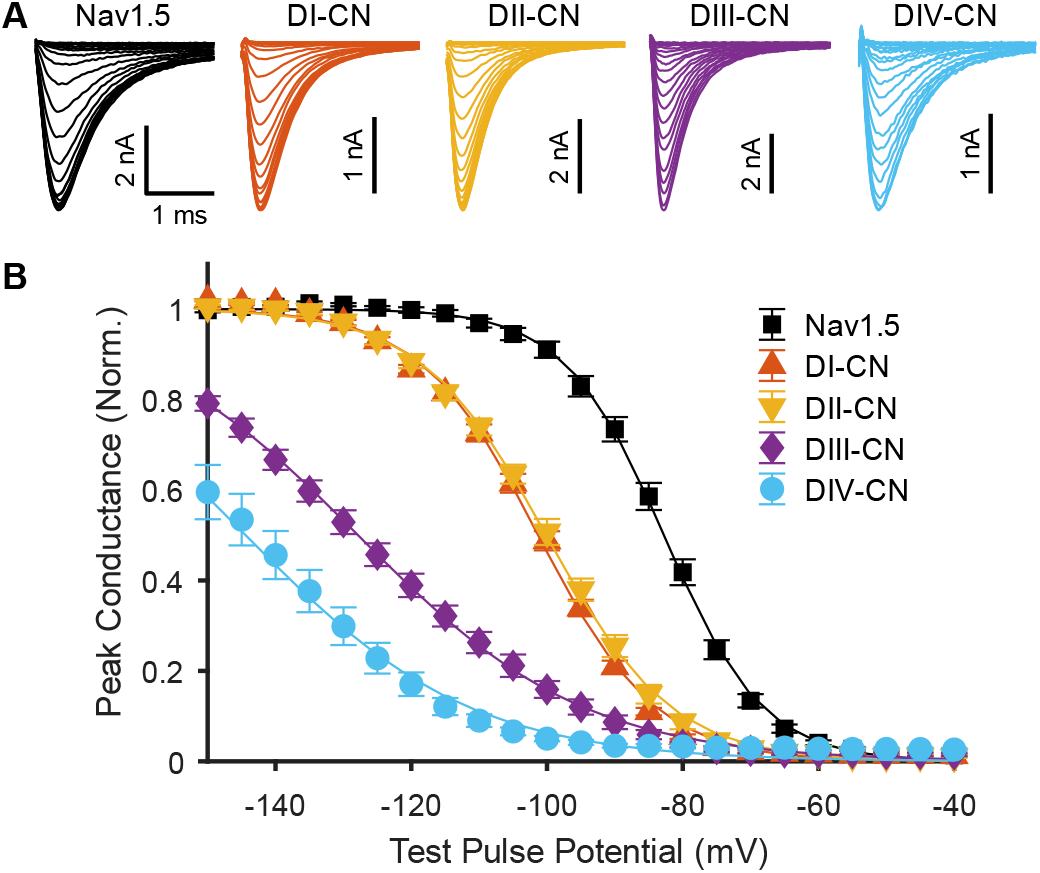
Changes in steady-state inactivation following domain neutralization in adult form of Nav1.5, mH1. (**A**) Representative traces of ionic currents corresponding to wild-type (WT) Nav1.5 (cell 20170311c3) and mutant channels (DI-CN, cell 20180301c6; DII-CN, cell 20180711c3; DIII-CN, cell 20180309c12; DIV-CN, cell 20190305c4) in response to a test pulse to −10 mV following conditioning pulses ranging from −160 to −30 mV. (**B**) Summary data corresponding to panel **A**, showing normalized peak current following the test pulse, as a function of the conditioning pulse voltage.

**Fig. S4.**
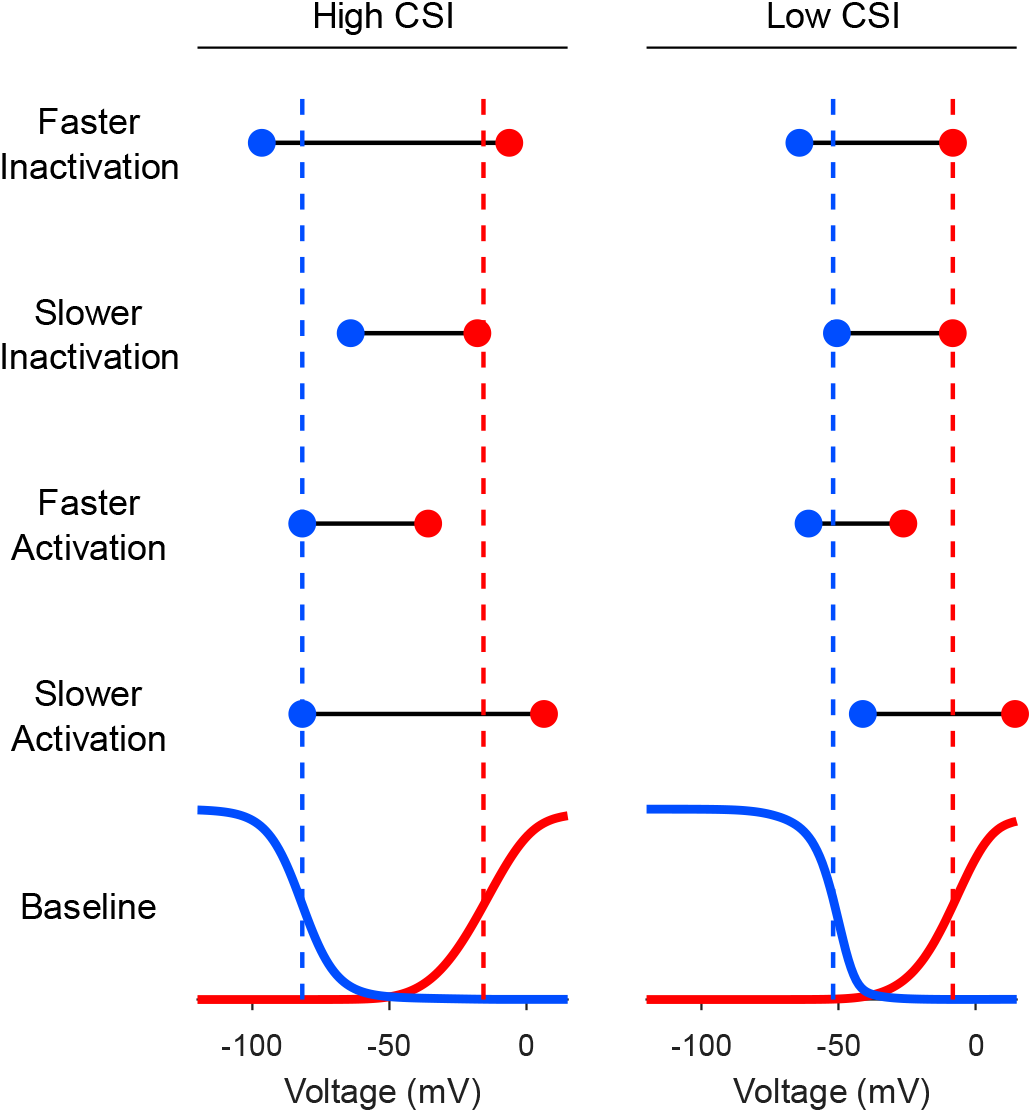
Comparison of models with high and low propensities for closed-state inactivation (CSI). High CSI refers to the reparametrized Nav1.5e model (Fig. 2A and Table S2). Low CSI refers to the same model, where inactivation has been biased towards open-state inactivation (see Methods). The GV (red) and SSI (blue) curves of these two models are depicted at the bottom (baseline). Above these curves are depicted the changes in the V_1/2_ of GV (red circle) and SSI (blue circle) following four different parameter modifications, with the dashed lines providing a reference to the baseline values: faster inactivation, α_4_ increased 10-fold; slower inactivation, α_4_ decreased 10-fold; faster activation, γ increased 10-fold; slower inactivation, γ decreased 10-fold.

**Table S1.** Boltzmann parameters for Nav1.5e.

| Nav1.5e | $V_{1/2}$ | k | n |
| --- | --- | --- | --- |
| WT<sup>†</sup> | $-16.8 \pm 0.5$ | $9.0 \pm 0.1$ | 51 |
| | $-82.0 \pm 0.5$ | $-7.2 \pm 0.1$ | 49 |
| DI-CN | $-48.1 \pm 0.6$ | $9.9 \pm 0.1$ | 24 |
| | $-100.9 \pm 0.5$ | $-8.7 \pm 0.1$ | 23 |
| DII-CN | $-22.6 \pm 0.5$ | $9.6 \pm 0.2$ | 26 |
| | $-96.5 \pm 0.8$ | $-9.2 \pm 0.2$ | 25 |
| DIII-CN | $-19.0 \pm 0.6$ | $10.3 \pm 0.2$ | 23 |
| | $-120.0 \pm 1.1$ | $-14.4 \pm 0.2$ | 23 |
| DIV-CN | $-7.4 \pm 1.3$ | $10.7 \pm 0.3$ | 14 |
| | $-131.1 \pm 2.6$ | $-13.7 \pm 1.5$ | 13 |
| + $\beta_1$ | $-14.8 \pm 0.7$ | $8.6 \pm 0.1$ | 14 |
| | $-73.5 \pm 1.0$ | $-6.1 \pm 0.3$ | 11 |
| + $\beta_3$ | $-13.8 \pm 0.8$ | $8.9 \pm 0.2$ | 18 |
| | $-74.5 \pm 0.6$ | $-6.2 \pm 0.2$ | 17 |
| DI*-VCF<sup>‡</sup> | $-65.4 \pm 2.3$ | $12.3 \pm 1.2$ | 13 |
| DII*-VCF | $-44.1 \pm 0.7$ | $22.2 \pm 0.6$ | 17 |
| DIII*-VCF | $-137.4 \pm 1.1$ | $15.9 \pm 0.6$ | 27 |
| DIV*-VCF | $-69.5 \pm 1.3$ | $13.2 \pm 0.5$ | 15 |

<sup>†</sup>Top row for each channel contains estimated Boltzmann parameters for peak activation (G/V) and bottom row contains parameters for steady-state inactivation (SSI). <sup>‡</sup>Parameters for VCF constructs are fits to the fluorescence-voltage (F-V) curve. Data report mean ± S.E.

**Table S2.** Parameters for Nav1.5e gating model.

| Rate constant | $k^*$ | $q$ | Cooperativity factor | |
| --- | --- | --- | --- | --- |
| $\alpha$ | 17,000 | -0.23 | $x_\alpha$ | 0.73 |
| $\beta$ | 830 | 1.4 | $x_\beta$ | 0.092 |
| $\alpha_4$ | 1,000 | -0.22 | $y_\alpha$ | 4.1 |
| $\beta_4$ | $3.0 \times 10^6$ | -0.43 | $y_\beta$ | 0.65 |
| $\alpha_{4o}$ | 1,800 | -0.7 | | |
| $\beta_{4o}$ | 74 | 1.7 | | |
| $\gamma$ | 10,000 | -3.0 | | |
| $\gamma_i$ | 51,000 | -5.2 | | |
| $\gamma_4$ | 17,000 | -6.2 | | |
| $\delta$ | 910 | -1.9 | | |
| $\delta_4$ | 690 | -2.2 | | |
| $\delta_i$ | 3,000 | -0.81 | | |
| $i$ | 25,000 | -0.013 | | |
| $r$ | 5,200 | 0.054 | | |
| $i_o$ | 11,000 | 0.11 | | |
| $r_o$ | 0.001 | 0 | | |

**Table S3.**
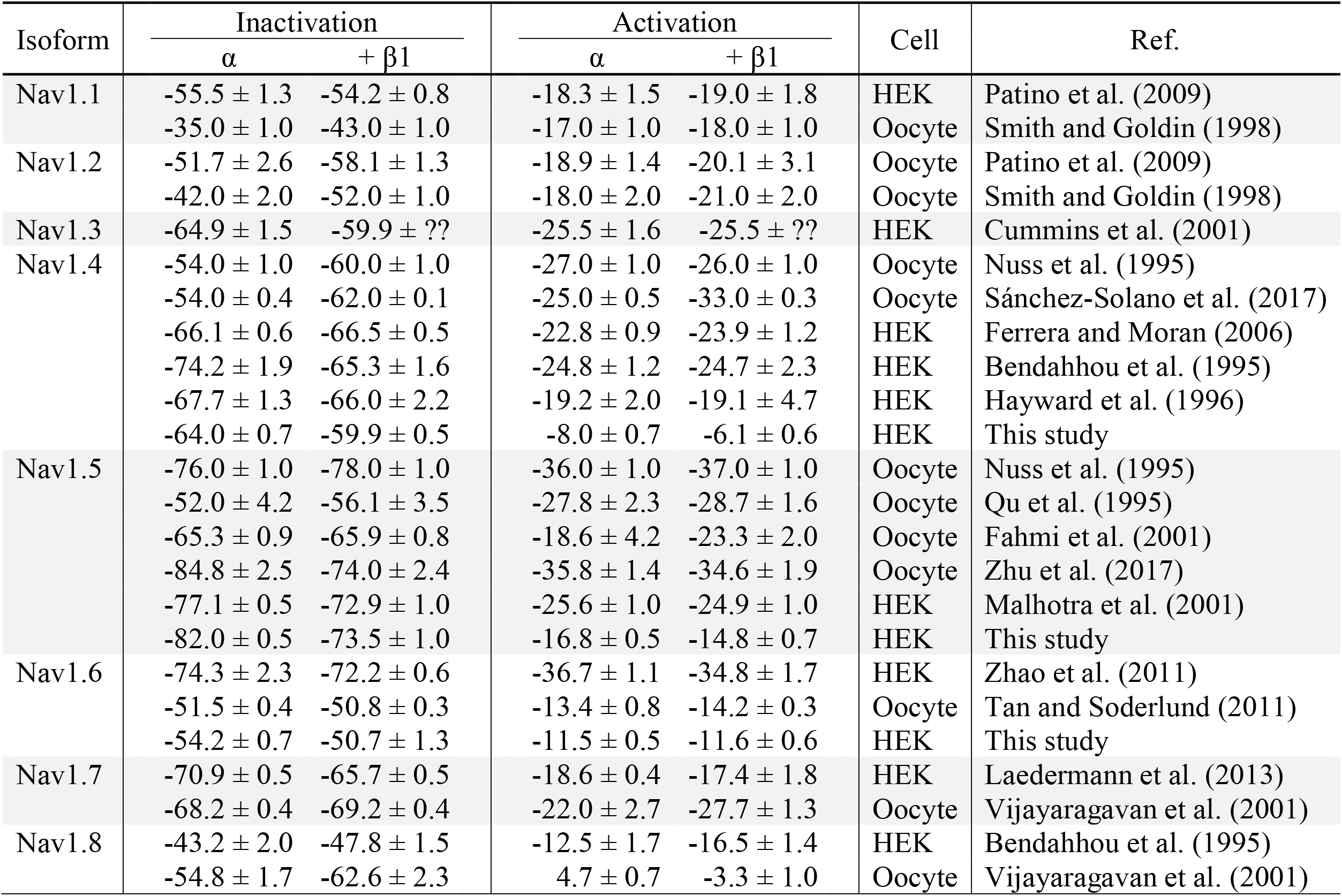
Summary for β1 effects on activation and inactivation.

